# Effectiveness of CRISPR-Cas in Sensitizing Bacterial Populations with Plasmid-Encoded Antimicrobial Resistance

**DOI:** 10.1101/2024.07.05.602127

**Authors:** Johannes Kippnich, Fabienne Benz, Hildegard Uecker, Franz Baumdicker

## Abstract

The spread of bacteria resistant to antibiotics poses a serious threat to human health. Genes that encode antibiotic resistance are often harbored on plasmids, extra-chromosomal DNA molecules found in bacteria. The emergence of multiresistance plasmids is particularly problematic and demands the development of new antibiotics and alternative strategies. CRISPR-Cas derived tools with their sequence specificity offer a promising new approach to combating antibiotic resistance. By introducing CRISPR-Cas encoding plasmids that target antibiotic resistance genes on plasmids, the susceptibility of bacteria to conventional antibiotics can be restored. However, genetic variation within bacterial populations can hinder the effectiveness of such CRISPR-Cas tools by allowing some mutant plasmids to evade CRISPR-mediated cleaving or gene silencing. In this study, we develop a model to test the effectiveness of CRISPR-Cas in sensitizing bacterial populations carrying resistance on non-transmissible plasmids and assess the success probability of a subsequent treatment with conventional antibiotics. abstract hgt We evaluate this probability according to the target interference mechanism, the copy number of the resistance-encoding plasmid, and its compatibility with the CRISPR-Cas encoding plasmid. Our results identify promising approaches to revert antibiotic resistance with CRISPR-Cas encoding plasmids: A DNA-cleaving CRISPR-Cas system on a plasmid incompatible with the targeted plasmid is most effective for low copy numbers, while for resistance plasmids with higher copy numbers gene silencing by CRISPR-Cas systems encoded on compatible plasmids is the superior solution.

## Introduction

The escalating crisis of antimicrobial resistance (AMR) poses a serious global health threat. Projections suggest that the number of deaths associated with AMR could rise to over 8 million per year over the next 25 years (Naghavi *et al*. 2024). Particularly in bacteria, the misuse and overuse of antibiotics has led to the rapid evolution of resistance (Schrader *et al*. 2020; Antimicrobial Resistance Collaborators 2022), resulting in a constant demand for novel effective antibiotics. While the number of multidrug-resistant pathogens increases, the rate of antibiotic discovery has declined in recent years (Luepke *et al*. 2017). Consequently, there is a pressing need for innovative strategies to treat infections with resistant bacteria.

AMR genes are often located on plasmids, which are self-replicating extrachromosomal DNA elements (Rozwandowicz *et al*. 2018; Castañeda-Barba *et al*. 2024). Plasmids self-regulate their replication through negative-feedback loops, and often exist in multiple copies within bacterial cells (Ramiro-Martínez *et al*. 2024). Higher plasmid copy numbers increase cellular levels of AMR-conferring enzymes through gene dosage (Friehs 2004; Hernandez-Beltran *et al*. 2024) and the occurrence of novel mutations in plasmid-carried genes (San Millan *et al*. 2016; Santer and Uecker 2020). If two plasmids encode very similar partition and/or replication functions, their regulatory systems interfere with each other, leading to the loss of one of the two plasmids over multiple generations, which thus makes them incompatible (Ebersbach and Gerdes 2005; Bouet *et al*. 2007). Some plasmids encode a dedicated transfer machinery, allowing them to spread horizontally to neighboring bacterial cells via conjugation. Such self-transmissible plasmids largely drive the spread of multidrug resistance in bacterial pathogens (von Wintersdorff *et al*. 2016; Partridge *et al*. 2018). For a more detailed guide to plasmid biology – targeted specifically also at modelers –, see Dewan and Uecker (2023).

One promising alternative to developing new antibiotics is the removal of plasmid-encoded bacterial resistance by employing CRISPR-Cas derived tools, rendering bacteria sensitive to conventional antibiotics (Buckner *et al*. 2018). CRISPR-Cas are sequence-specific immune systems that protect their bacterial hosts from infecting mobile genetic elements such as bacteriophages and plasmids. Sequence specificity is achieved by CRISPR RNAs (crRNAs) that guide the effector proteins to their complementary sequences for target interference. The molecular mechanisms by which CRISPR-Cas systems recognize and interfere with their targets vary greatly, but generally, they provide host immunity by cleaving of or binding to a target nucleic acid (Nussenzweig and Marraffini 2020). CRISPR-Cas systems have been exploited extensively to develop versatile genetic engineering tools (Benz *et al*. 2025; Selle and Barrangou 2015; Adli 2018) that can be directed to a specific genomic locus by providing crRNAs base-pairing with the target sequence. CRISPR-Cas tools have also been developed into antimicrobials: Cleaving CRISPR-Cas tools directed to the bacterial chromosome lead to sequence-specific killing (Bikard *et al*. 2014; Citorik *et al*. 2014) and when directed towards antibiotic-resistance plasmids, bacteria can be sensitized to antibiotics (Citorik *et al*. 2014; Yosef *et al*. 2015; Wang *et al*. 2019; Zhou *et al*. 2023). As an alternative to removing plasmids by target cleaving, effectors that result in transcriptional repression of target genes can be used to silence AMR (Yao *et al*. 2022; Mayorga-Ramos *et al*. 2023).

For CRISPR-Cas-based antimicrobials to be effective, the effector needs to be delivered to the entire target population. Due to the high infectivity of bacteriophages, phage-mediated delivery has been tested extensively (Park *et al*. 2017; Lam *et al*. 2021; Brödel *et al*. 2022; Gencay *et al*. 2024), and plasmid transfer has been deemed not efficient enough (Citorik *et al*. 2014). However, more recent *in vivo* studies highlight the potential of conjugative plasmids and their engineered derivates as potent delivery devices (Ronda *et al*. 2019; Rodrigues *et al*. 2019; Neil *et al*. 2020; Jin *et al*. 2022; Walker-Sünderhauf *et al*. 2023).

Nevertheless, just as to antibiotics, bacteria can evolve resistance to such CRISPR-Cas antimicrobials by mutation of the target sequence, or the CRISPR-Cas tool itself may acquire mutations, rendering it ineffective (Mayorga-Ramos *et al*. 2023). Sensitizing the bacterial population followed by antibiotic treatment strongly selects for mutations that evade CRISPR-Cas targeting and consequently result in a rise of antimicrobial resistant bacteria, provided the escape mutation has not compromised the function of the resistance gene.

Although experimental proof-of-concept studies showed promising results (Yosef *et al*. 2015; Wang *et al*. 2019; Zhou *et al*. 2023; Walker-Sünderhauf *et al*. 2023), the specific conditions under which a CRISPR-Cas sensitizing strategy can be effective, even when mutations allow CRISPR-Cas evasion, remain uncertain. Mathematical models have been crucial to understand the evolutionary dynamics of plasmids, but models that explain the dynamics of plasmids targeted by CRISPR-Cas (Mamontov *et al*. 2022) and the effectiveness of such strategies are scarce.

Here, we develop a comprehensive mathematical model to explore the effectiveness of CRISPR-Cas tools in removing plasmid-encoded AMR. We utilize Moran models and multitype branching processes and consider different design options for the CRISPR-Cas tool. We consider the CRISPR-Cas-encoding plasmid to be either compatible or incompatible with the AMR-conferring plasmid, and the CRISPR-Cas effectors to act by either cleaving or by binding and transcriptionally silencing their target (the AMR gene). Our models suggest that the optimal strategy for targeting AMR plasmids depends on their copy number. If the copy number of the AMR plasmid is below a certain threshold, a plasmid that is incompatible with the targeted AMR plasmid and delivers a target-cleaving CRISPR-Cas tool is most effective for sensitizing. If the copy number of the targeted AMR plasmid is above this threshold, delivery of a gene-silencing CRISPR-Cas tool by a compatible plasmid is the superior strategy under most circumstances.

## The model

purpose The purpose of our model is to evaluate the effectiveness of CRISPR-Cas in sensitizing bacterial populations and to determine the probability of their eradication through subsequent antibiotic treatment.

model start We consider a population of bacterial cells that each carry a combination of three plasmid types:

### pAMR

Wild-type (WT) AMR plasmid that provides the host cell with resistance to a given antibiotic.

### pAMRmut

Mutated AMR plasmid that evades CRISPR-Cas targeting via a mutation in the target gene and – assuming a worst-case scenario – provides the host with the same level of AMR during treatment as the WT AMR plasmid.

### pCRISPR

Plasmids that encode a CRISPR-Cas tool programmed to target the AMR gene of pAMR. We consider two types of pCRISPR that differ in their mode of action:

1. **pCleaving:** Introducing double-strand DNA breaks (as for example type I and II CRISPR-Cas systems), and
2. **pSilencing:** Silencing the AMR gene of pAMR by transcriptional repression (as for example type IV CRISPR-Cas).

The combination of these plasmids within a cell determines the birth and death rates of the cell (in the following referred to as the fitness of a cell) and defines the cell type. temporal The model operates in continuous time, where cell division and death occur after exponentially distributed waiting times. HGT first mention For simplicity, horizontal transfer is excluded for all three plasmid types in the model; its role is addressed in the discussion.

spatio temporal We focus on the population genetics processes and simplify the ecology as much as possible. Specifically, we assume a well-mixed environment without spatial structure, ignore all ecological interactions between cells such as competition for resources during antibiotic treatment, and assume the antibiotic concentration to be constant.

### Process overview

The model consists of three steps, illustrated in Fig. 1: Concept Figure line

**Fig. 1.**
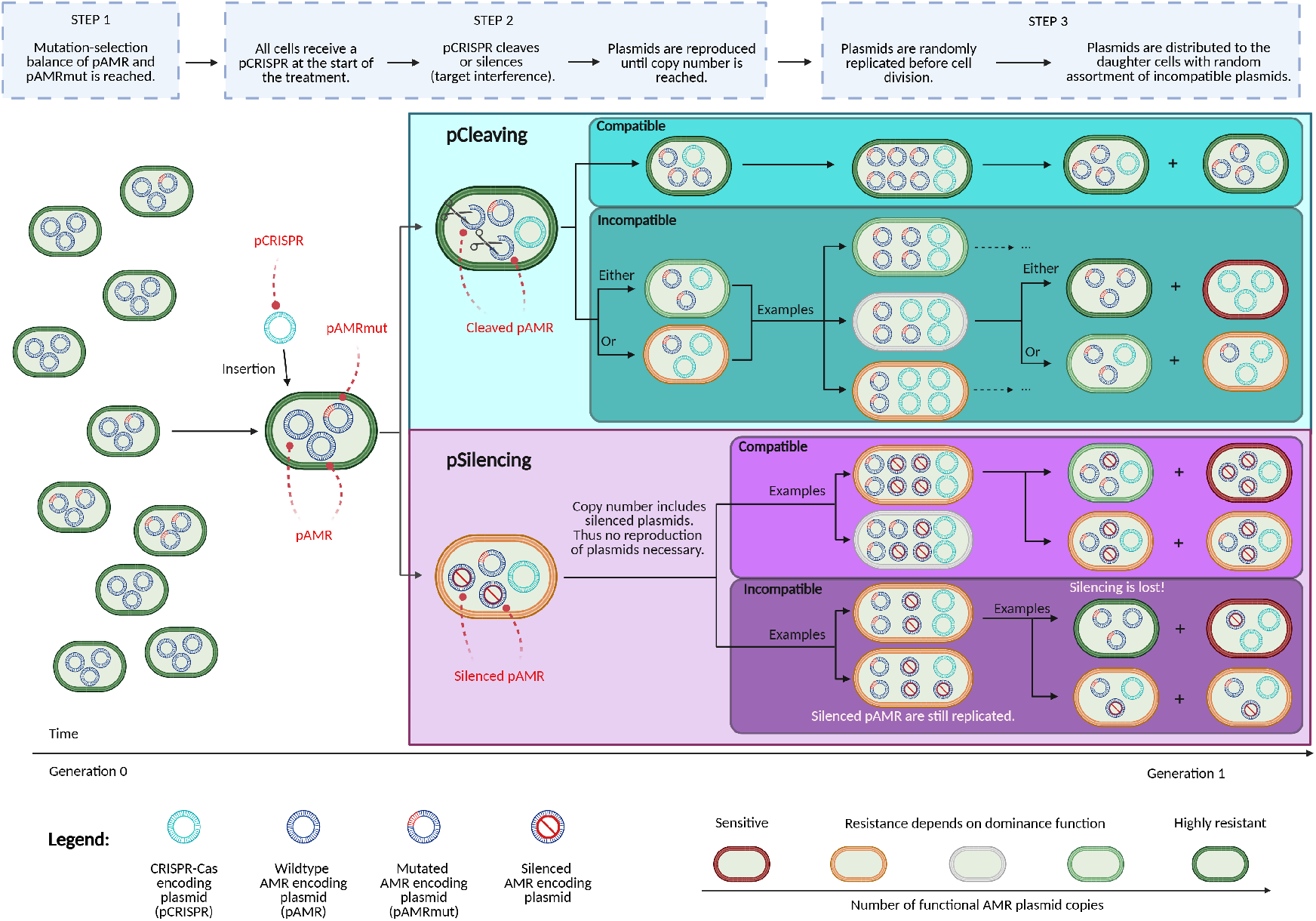
Sensitizing concept and the interplay of pCRISPR and the AMR plasmid replication and segregation. All cells in the population of pathogens carry a multicopy antimicrobial resistance (AMR) plasmid (pAMR), where some copies (pAMRmut) might carry a mutation that makes them immune to a CRISPR-Cas-encoding plasmid (pCRISPR). pCRISPR can have a cleaving (pCleaving) or transcriptional repression (pSilencing) mode of action, in both cases directly targeting the AMR gene on the plasmid (see step 1 in model description). At the beginning of the CRISPR-Cas treatment (step 2), each cell receives a copy of pCRISPR, which acts immediately: For pCleaving all wild-type pAMR copies are cleaved, and pAMRmut copies can undergo replication until the plasmid copy number is reached. Conversely, when silencing takes place, there is no imperative for reproduction. In the following step 3, cell division and death proceed under antibiotic treatment in accordance with individual fitness, determined by the quantity of non-silenced AMR plasmids retained within the cells. Throughout each cell division, the plasmids are replicated and equally distributed to the progeny cells with random assortment of plasmid variants. This results in different combinations of mother-daughter cells, introducing variability in the inheritance of plasmids during cell division. Created in BioRender. Baumdicker, F. (2025) https://BioRender.com/d97v528

#### Step 1: Standing genetic variation prior to any treatment (*t <* 0)

In this initial stage before the introduction of pCRISPR and antibiotic treatment, cells can harbor pAMR and/or pAMRmut. We assume that the CRISPR-Cas evading mutation in pAMRmut carries a small cost, and that the population is in mutation-selection balance which determines the initial distribution of pAMRmut.

#### Step 2: Sensitizing via pCRISPR introduction (*t* = 0)

At the start of this treatment step, each cell acquires exactly one pCRISPR in addition to the existing pAMR and pAMRmut. The effect of pCRISPR on pAMR is assumed to be instantaneous, i.e. the WT pAMR copies are immediately cleaved or silenced. Note that silencing is reversible if pSilencing is lost from a cell in later generations.

#### Step 3: Treating the population with antibiotics (t > 0)

The antibiotic treatment starts as soon as each cell has received pCRISPR. Cells without pAMRmut are sensitive to antibiotic treatment, as all pAMR are cleaved or silenced. The survival probability of cell lineages that originate from cells with either preexisting or*de novo* acquired pAMRmut copies depends on demographic stochasticity, the initial number of pAMRmut copies, and the mode of action and compatibility type of pCRISPR.

With this model, we determine the extinction probability of the bacterial population, or conversely, the potential for evolutionary rescue through mutations in pAMR. A description of all relevant parameters is provided in Table 1. Table 1 reference See Supplementary Note 1ODD protocoll verweis for a description of our model using the ODD (Overview, Design concepts, Details) protocol (Grimm *et al*. 2020). In the following, we detail considerations for steps 1 - 3.

**Table 1.** Description of all relevant parameters in the model(s).

|  |  |
| --- | --- |
| N | Number of cells before pCRISPR introduction ($t \leq 0$) |
| n | Copy number of pAMR and pAMRmut |
| v | Probability for a pAMR to mutate to pAMRmut during replication |
| s | Cost of the mutation in the absence of antibiotics ($t < 0$) |
| $\lambda_k^{(n)}$ | Division rate of cells with $k$ functional AMR plasmids (pAMRmut or non-silenced pAMR) and copy number $n$ ($\lambda_{\min}^{(n)} = \lambda_0^{(n)}, \lambda_{\max}^{(n)} = \lambda_n^{(n)}$) |
| $\mu_k^{(n)}$ | Death rate of cells with $k$ functional AMR plasmids and copy number $n$ |
| $R_k^{(n)}$ | Fitness/Birth-death ratio<br>($= \lambda_k^{(n)} / \mu_k^{(n)}$) |

#### Step 1: Standing genetic variation prior to any treatment

Some cells in the bacterial population may already contain pAMRmut or a mixture of pAMR and pAMRmut copies prior to the introduction of pCRISPR. To simulate the frequency of pAMRmut at the time of pCRISPR introduction, we utilize a multitype Moran model, as developed by Santer and Uecker (2020). This model assumes a constant population of *N* bacterial cells carrying pAMR copies, where each cell contains exactly *n* plasmids. plasmid free cells In particular, we do not take plasmid-free cells into account. In reality, plasmid-free cells may emerge as a result of segregational loss and rise in frequency due to the fitness cost of plasmid carriage in the absence of antibiotics (Nicoloff *et al*. 2024). Nevertheless, plasmid-free cells are effectively killed by the antibiotic treatment irrespective of the pCRISPR variant, and plasmid transfer within the population that could reintroduce plasmids into plasmid-free cells is not modeled. Therefore, ignoring plasmid-free cells does not affect the conclusions here.

Initially, all plasmids are pAMR (i.e., # pAMR = *n* and # pAMRmut = 0). In each Moran model time step, one cell is randomly selected to die, while another one is randomly selected to divide. This mother cell produces two daughter cells, replacing itself and the deceased cell to maintain a constant population size. During each cell division, each plasmid is replicated exactly once resulting in a total number of 2*n* copies, which are then distributed to the daughter cells with random assortment of plasmid variants. (We provide details on plasmid replication and segregation in the section ‘Copy number and plasmid incompatibility’ below.) At each plasmid replication event, the CRISPR-Cas target sequence in pAMR mutates with probability *v*, thereby transitioning from pAMR to pAMRmut. We assume that prior to the introduction of pCRISPR the mutation in pAMRmut has a small cost for the host cell and assume that this selective cost increases linearly with the number of pAMRmut copies up to a maximum cost *s*. The costs determine the probability that a cell is selected for replication in the Moran model such that cells with *j* pAMRmut copies have a reduced weight of 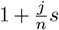 (*s* is negative). Consequently, in the long run, the frequency of pAMRmut copies will evolve towards mutation-selection balance, and pAMRmut will be present at a low frequency when pCRISPR is introduced in the next step (Fig. 1; left). SGV mutational input The standing genetic variation (SGV) in natural populations will, of course, differ from our model. However, as we will see below, the essential properties under which our overall conclusions are expected to hold are generally plausible: pAMRmut preexists in different copy numbers in a small fraction of the population, and low pAMRmut copy numbers are more common than high copy numbers if *n* is large.

#### Step 2: pCRISPR introduction

Upon pCRISPR introduction (t=0), we assume that each cell receives exactly one pCRISPR copy (Fig. 1; center-left). cell which not gain crispr In reality, some cells may remain pCRISPR-free and retain AMR due to imperfect pCRISPR distribution. However, while such cells would decrease the extinction probability of the population, this effect would be identical for all pCRISPR variants and therefore does not affect their relative comparison, which is the focus of our study.

The interference activity is effective immediately upon delivery and the model makes the following assumptions for each mode of action of pCRISPR:

##### pCleaving

All pAMR copies within a given cell are cleaved and thus lost before the next cell division. In contrast, pAMRmut copies are protected by their mutation and replicate immediately according to their replication mechanism.

##### pSilencing

pAMR remains in the cell, but the expression of the AMR gene is silenced. Therefore, pAMR does no longer provide antibiotic resistance. pAMRmut is protected from transcriptional repression. The replication of pAMR and pAMRmut is not affected by target silencing. If all pSilencing copies are lost from a cell in later generations due to plasmid segregation, the pAMR copies become functional and provide resistance again.

#### Step 3: Treatment with antibiotics

The antibiotic treatment starts as soon as each cell has received pCRISPR. Treatment selects for pAMRmut which carries a functional resistance gene.

To simulate the bacterial population dynamics, we employ a birth-death process similar to the one described in Santer and Uecker (2020). Here, for each cell, the division and death rate depends on the total number *k* ∈ {0, …, *n*} of functional resistance plasmids, i.e. pAMRmut and non-silenced pAMR copies within the cell (Fig. 1; right) concept figure II. We deliberately ignore any differential fitness costs associated with the total plasmid load to focus on a comparison of the inherent differences between pCRISPR design choices with respect to different copy numbers. birth and death rates Accordingly, there are a total of *n* + 1 cell types with different birth rates 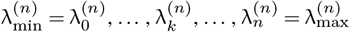, where *k* is the sum of all functional AMR plasmid copies. The death rates 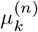 are defined analogously.

mu = 1lambda and mu The extinction probability of the birth-death process depends solely on the probability of each cell to divide or die – specifically, the ratio of the birth and death rates 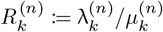. birth-death ratio We refer to *R*_*k*_ as the fitness of a cell in the following. We further define 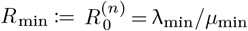 and 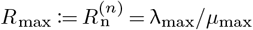. In the following, we assume 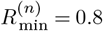 (if not stated otherwise), which results in a population decline of completely sensitized cells by 50% within six steps in the discrete-time process in which cells either divide or die in each step or – going back to the continuous-time birth death process – within four bacterial generations (measured as interdivision times 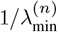).

Furthermore, we assume that for cells with the maximum amount of functional AMR plasmids, 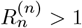, since the bacterial WT population would face certain extinction otherwise, even without any sensitization. This can be easily seen when looking at the Malthusian fitness, that is the difference 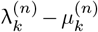, which for *k* = *n* results in a growing population only when 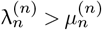.

For cells with *k* ∈ {1, …, *n* − 1}, it is also natural to assume that for the same copy number *n* more functional AMR plasmids lead to a higher replication rate λ_*k*_ and/or a lower death rate *µ*_*k*_, such that 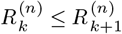. This inspires the following fitness functions, which we will consider in detail (see Supplementary Fig. S4 for an interpretation in the context of bacteriostatic vs bactericidal antibiotics):

##### *Linear* fitness

The fitness increases linearly with the number of functional AMR plasmids, i.e. 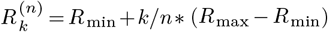.

##### *Dominant* AMR plasmids

One functional AMR plasmid is sufficient to achieve maximum fitness for the cell, i.e. 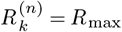 for 1 ≤ *k* ≤ *n*.

##### *Recessive* AMR plasmids

The cell needs the maximum amount of functional AMR plasmids, otherwise there is no fitness effect of the AMR plasmids, i.e. 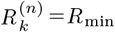 for 0 ≤ *k* ≤ *n* − 1.

definition dominance Note that we use the terms ‘dominant’ and ‘recessive’ here since the functions describe the effect of the relative amount of functional AMR plasmids among the *n* incompatible plasmids on the phenotype. However, unlike in the usual population genetic context, we are not considering dominance of one allele over another.

We assume in the following that the maximum number of functional AMR plasmids, *n*, always provides the same maximum cell fitness, irrespective of the copy number *n*. When considering different resistance genes on different plasmids, results can be read as ‘success probabilities, given a certain copy number and maximum cell fitness’. If we think of the same resistance gene on different plasmids, the assumption of a fitness that does not depend on *n* is inconsistent with resistance mechanisms such as antibiotic-degrading enzymes for which a higher number of AMR plasmids results in a higher level of AMR (Stougaard *et al*. 1979; San Millan *et al*. 2016; Santos-Lopez *et al*. 2017). However, it allows for a direct comparison of the impact of the copy number on the success of sensitizing CRISPR-Cas treatment without the confounding effects of gene dosage.

### Copy number and plasmid incompatibility

Except for the CRISPR-Cas evading mutation in the AMR gene, pAMR and pAMRmut are isogenic and rely on the same copy number control system and are thus incompatible. For pCRISPR, we consider two scenarios: one in which pCRISPR shares the same incompatibility group as pAMR/pAMRmut (hereafter referred to as incompatible pCRISPR), and one in which it belongs to a different incompatibility group (hereafter referred to as compatible pCRISPR). For mathematical convenience, we assume that each cell contains exactly *n* copies of the incompatibility group of pAMR/pAMRmut, unless they have all been cleaved, although in reality there is some variation between cells (Shao *et al*. 2021).

If we denote the number of pAMR copies within a cell as #pAMR (and accordingly for the other plasmids), this leads, depending on the (in-)compatibility between pCRISPR and the resistance plasmid, to:

#### Incompatible pCRISPR

#pAMR + #pAMRmut + #pCRISPR = *n* and

#### Compatible pCRISPR

#pAMR + #pAMRmut = *n*.

Whether pCRISPR is compatible or incompatible with pAMR/pAMRmut may have drastic consequences for the treatment outcome. If incompatible, the replication of pCRISPR interferes with the replication and segregation of pAMR and pAMRmut because it is perceived as the same plasmid by the regulatory system and thus over time ultimately results in the loss of either pCRISPR or pAMR and pAMRmut. The details of this effect are outlined in the next section. For compatible pCRISPR, there is no interference with the replication of pAMR and pAMRmut and thus compatible pCRISPR is assumed to never be lost. Note that the copy number of compatible pCRISPR therefore does not influence our model as long as it is stably maintained.

### Plasmid replication and segregation during cell division

Each time a cell divides, all plasmids incompatible with pAMR, including the pAMR copies themselves, replicate to 2*n* copies (Fig. 1; center-right). In our calculations, we consider two cases for the plasmid replication system: one in which the system perfectly remembers which plasmid has already been copied, and one where the system is memoryless, as described in Novick and Hoppensteadt (1978) and Nordström (2006): If the plasmids replicate *regularly*, all plasmids are replicated exactly once. If the plasmids replicate *randomly*, a random plasmid is chosen for replication and then added to the pool until the copy number has been reached. The random replication model, where the “rich get richer”, is known in mathematics as the Pólya urn scheme (Pólya 1930). The frequency of the different plasmid types, pAMR, pAMRmut, and incompatible pCRISPR, among the replicated plasmid copies is then Dirichlet-multinomial (DirMult) distributed. In our framework, we only use regular replication to infer the standing genetic variation but compare the two alternative replication models during treatment. Our analysis shows that the deviations between results for the two replication models are minimal for linear fitness, even quantitatively, and qualitatively similar for dominant and recessive fitness, such that the overall findings hold for both replication models. For the results depicted in Figures 2 - 5, we use the random replication model. The results based on regular replications are depicted in Supplementary Figures S1 - S3. A more general comparison of the two replication models can be found in Santer *et al*. (2022) and Dewan and Uecker (2024).

**Fig. 2.**
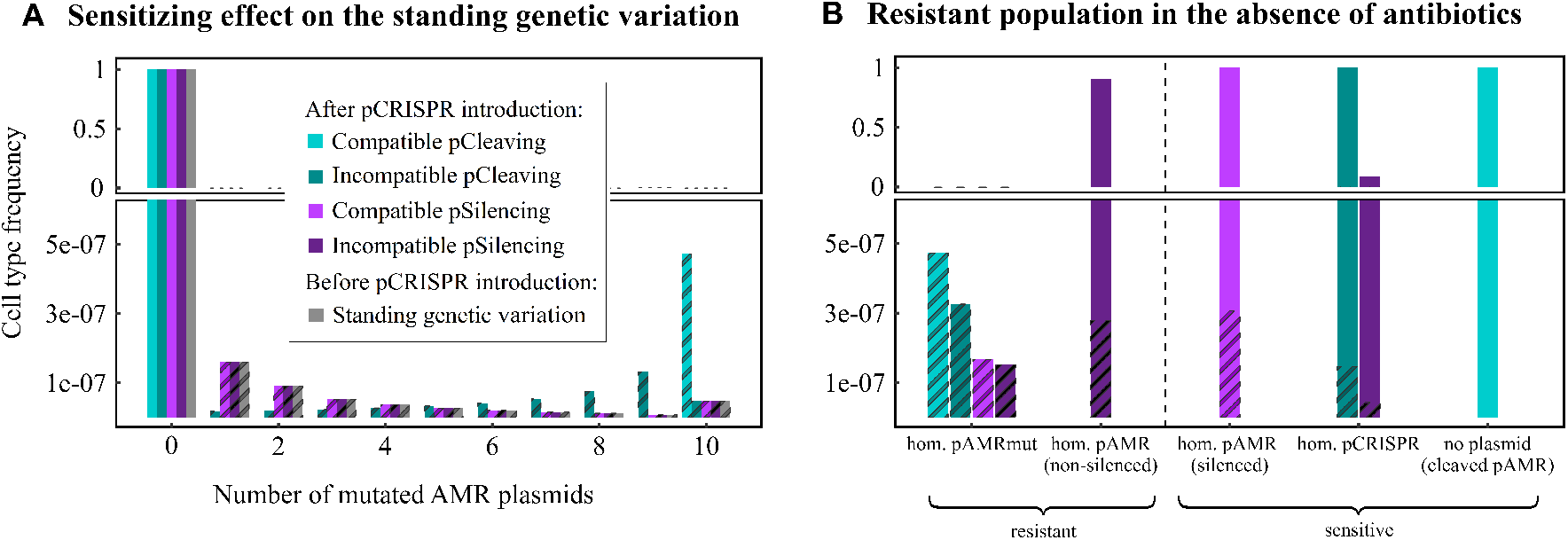
The standing genetic variation (SGV) before and after the introduction of pCRISPR to the population for copy number *n* = 10. (A) The frequency of cells carrying zero pAMRmut up to the maximum number of *n* = 10 copies. The remaining plasmids within the cells are either wild-type pAMR or pCRISPR copies. Cells with at least one pAMRmut copy are represented with striped bars. The distribution before the introduction of pCRISPR to the population is depicted in grey. The colored bars show the frequency of all cell types immediately after the introduction of one pCRISPR copy to each cell. These frequencies depend strongly on the pCRISPR variant. Compatible pCleaving (light turquoise) transforms all heterozygous cells into homozygous mutant cells, as all pAMR are cleaved and replaced by pAMRmut, as long as there is at least one pAMRmut copy present in the cell. Similarly, incompatible pCleaving (dark turquoise) increases the frequency of cells with high numbers of pAMRmut copies, as pAMRmut will be replicated alongside pCleaving after the pAMR have been cleaved. On the other hand, pSilencing (purple) does not change the plasmid composition, keeping a distribution where most heterozygous cells have a low number of pAMRmut copies. (B) Cell type frequencies many generations after the introduction of pCRISPR. Starting from the population shown in A and ignoring the effects of *de novo* mutations and antibiotic treatment all cells will become homozygous for a specific randomly chosen plasmid type over time due to random plasmid segregation. The descendants of cells with at least one pAMRmut copy (striped bars in A) are depicted with stripes. The resulting cell types are either resistant or sensitized. The frequency on the y-axis is shown once from 0 to 6 ∗ 10^−7^ and once from 0 to 1 to account for the different scales. Parameters: *v* = 10^−8^, *n* = 10, *s* = −0.1.

After plasmid replication, the cell divides, with each daughter cell receiving exactly *n* plasmid copies. The copies of incompatible plasmids are distributed hypergeometrically (see Novick and Hoppensteadt (1978) for more details on this model). plasmid replication While these assumptions simplify the complex biological process of plasmid segregation, they enable a mathematically tractable formulation.

In summary, for a mother cell with *j* pAMRmut copies undergoing cell division, this implies that after random plasmid replication to 2*n* plasmid copies, the number of newly generated pAMRmut copies *j*_*d*_ is DirMult (*n*, (*j, n* − *j*)) distributed. The number of pAMRmut copies in the resulting two daughter cells *j*_1_ + *j*_2_ = *j* + *j*_*d*_ are then distributed such that *j*_1_ ∼ Hypergeom(2*n, j* + *j*_d_, *n*).

## Analysis

Our primary goal is to calculate the extinction probability of a bacterial population undergoing antibiotic treatment after receiving one of the four considered pCRISPR variants: compatible pCleaving, compatible pSilencing, incompatible pCleaving, and incompatible pSilencing.

To compute this probability, we first need to determine the frequency of pAMRmut copies in our Moran model when pCRISPR is introduced into the cells (step 1). Subsequently, we can calculate the frequency of each cell type right after pCRISPR introduction, which takes immediate effect according to its type and incompatibility group (step 2). Then, we can assess the probability of population extinction, denoted as *P*_ext_ (step 3).

### Standing genetic variation before pCRISPR introduction

The multitype Moran model with a constant population size, as described in step 1 of the previous section, has two key factors that affect the frequency of cells carrying a specific number of pAMRmut copies: the probability *v* of mutation occurrence and the reduction in fitness *s* for cells carrying pAMRmut copies. We assume that the population is in mutation-selection balance when the CRISPR-Cas plasmid is introduced and describe the composition of the standing genetic variation by the deterministic equilibrium. This equilibrium is computed by integrating nonlinear systems, following the calculations and code outlined in Appendix C of Santer and Uecker (2020), where for simplicity at most one mutation can occur per cell division of cells that do not yet carry pAMRmut copies. We denote by *f* ^(SGV)^(*j*) = *f* ^(SGV)^(*n* − *j, j*, 0) the fraction of cells that carry *j* copies of pAMRmut and *n* − *j* copies of pAMR immediately before introduction of pCRISPR.

### Introduction of pCRISPR

For compatible and incompatible pSilencing, the number of pAMR and pAMRmut copies remains the same after introduction as before, but all pAMR are silenced. In the case of compatible pCleaving, each cell with initially *j* pAMRmut copies is transformed into a cell with *n* pAMRmut copies. If pCleaving is incompatible, one pCleaving copy and *j* pAMRmut copies will compete for the remaining *n* − *j* − 1 spots. In the case of *randomly* replicating plasmids, we use the Pólya urn scheme to fill the remaining places as described in the previous section. For *regularly* replicating plasmids, we repeatedly double the remaining *j* + 1 plasmids unless their number would exceed *n*. The remaining spots are filled by randomly picking plasmids without replacement to replicate.

The resulting distribution of the fraction of cells immediately after the introduction of pCRISPR will be denoted by *f* ^(CRISPR)^(*i, j, c*), where *i* is the number of pAMR copies, *j* represents the number of pAMRmut copies, and *c* denotes the number of pCRISPR copies.

For compatible pCRISPR we have *i* + *j* = *n*, while for incompatible pCRISPR usually *i* + *j* + *c* = *n* holds. Note that introducing an incompatible pCRISPR will lead to the total number of plasmids within a cell exceeding *n*, when the cell was homozygous for pAMRmut or when using incompatible pSilencing. In these cases, this state will persist until the next cell division, during which pCRISPR, pAMR and pAMRmut will replicate so that there is a total of 2*n* of them. For random replication, this process is straightforward, whereas for regular replication, we simply assume that one random plasmid is not replicated. Subsequently, the plasmids are distributed to the daughter cells, each receiving *n* plasmids.

### Population extinction after pCRISPR introduction during antibiotic treatment

We are now interested in the probability that, once the CRISPR-Cas tool has been added for the purpose of sensitization, the entire bacterial population can be eradicated by antibiotics. How can the population escape from extinction? Irrespective of which pCRISPR variant is used, cells that carried a pAMRmut copy at the time of CRISPR-Cas treatment are not (fully) sensitized and can thus leave a growing lineage of progeny. For pCleaving, this is the only way in which the population can survive. For pSilencing, there are additional pathways to population rescue. If pSilencing is incompatible with pAMR, plasmid replication and segregation can lead to loss of pSilencing from cells, in which case the silenced AMR gene becomes functional again. Moreover, irrespective of the compatibility of pSilencing, new mutations from pAMR to pAMRmut can occur during the population decline following the introduction of pCRISPR. These mutations will be referred to as *de novo* mutations henceforth. Importantly, the presence of cells with functional resistance genes does not automatically lead to treatment failure since the lineage may still go extinct due to demographic stochasticity.

To account for this potential stochastic loss of (partially) resistant cells, we start by calculating the probability for each single cell to not produce a long-term lineage of offspring after the immediate effect of pCRISPR introduction, which will be a key building block of the analysis given further below. Since once a cell carries a pAMRmut copy, it is much more likely that the cell line survives thanks to that copy than through a new mutation occurring in future generations, we ignore the possibility of new mutations. Since the single-cell extinction probability is the same for cells of the same type, we associate each cell with its respective type, denoted as (*i, j, c*) as above. We then denote the single-cell extinction probability by *p* _ext_(*i, j, c*).

The calculation of *p* _ext_(*i, j, c*) can be done by using the independence of different branches in the birth-death process. In detail, this means that for a single cell the probability of extinction is the probability of immediate extinction, in addition to the probability of both lineages founded by the daughter cells going extinct after a possible cell division. For example, for *c* = 0 in the incompatible case and for any *c* in the compatible case, this simplifies to:

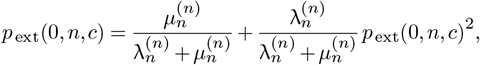

which yields 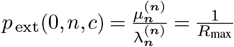, which is the known result for the extinction probability of a single-type birth-death process (Allen 2015, page 7).

Given a specific copy number *n* and considering all options for *i, j* and *c*, we get a total of *n* + 1 quadratic equations for compatible pCRISPR and 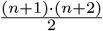 quadratic equations for incompatible pCRISPR. Due to the high number of equations to solve, we resort to numerical solutions. These solutions are based on properties of the probability generating function of multitype branching processes, as outlined in Appendix A of Santer and Uecker (2020).

Note that within this approximation (ignoring new mutations), *p*_ext_(*n*, 0, *c*) = 1 for compatible pSilencing, since all resistance plasmids remain silenced indefinitely. By contrast, loss of incompatible pSilencing from cells is rather likely, and it is therefore intuitive to expect (and confirmed in the Results section) that *p*_ext_(*n*, 0, *c*) is rather small, even in the absence of new mutations. Since this renders incompatible pSilencing ineffective, we will consider only compatible pSilencing, alongside both variants of pCleaving, in the following analysis of the total extinction probability of the bacterial population.

To calculate the total extinction probability, we then only have to consider the escape from extinction through either existing pAMRmut in the standing genetic variation or pAMRmut occurring through *de novo* mutations during population decline. We thus have to calculate the product of the extinction probabilities

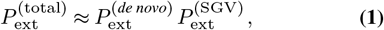

where 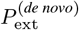 is the probability that no new mutations rescue the population from extinction and 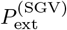 is the probability that none of the preexisting mutations allow the population to survive treatment.

To calculate the extinction probability 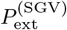, we multiply the extinction probabilities of all cell lineages:

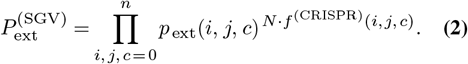

We now turn to 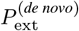, where only *de novo* mutations are taken into account. We first note that for pCleaving 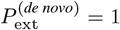, since there are no pAMR copies left after pCRISPR introduction and *de novo* mutations can thus not occur. To calculate the probability that *de novo* mutations do not lead to population rescue for compatible pSilencing, we focus on cells that were initially homozygous for pAMR and their progeny (cells that initially carry pAMRmut are covered by 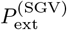). Because we are only interested in the total number of new mutations (and hence the number of cell divisions) until extinction of these cells and not their timing, we can consider discrete cell generations. In this discrete-generation process, a cell divides into two daughter cells with probability 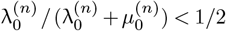 ; otherwise, it is dead. Assuming a deterministic decay of the cells carrying only pAMR, the number of cells therefore decreases by a factor 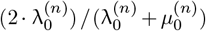 per generation. We can approximate the number of replications of pAMR during these cell divisions until population extinction by

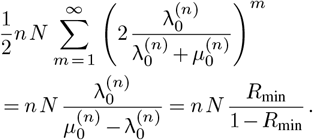

The factor 1*/*2 is due to only half of the plasmids of a new generation being freshly replicated, while the other half originates from the mother cells.

The total number of *de novo* mutations *M* is then distributed as Bin 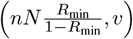, which can be approximated due to the Poisson limit theorem by Poi 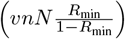 since the mutation probability *v* is small and the initial population size *N* is very large.

Once a cell has acquired a mutation, we can again ignore further mutations in the progeny. Overall, this results in the following extinction probability arising from *de novo* mutations only

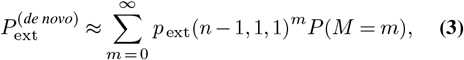

where we assume that only one *de novo* mutation occurs during a cell division and ignore the possibility that for a mutation that occurs in an early replication event, the mutant plasmid may be copied again in the *random* replication model (for a treatment of this possibility in a different context, see Dewan and Uecker 2024). Using the Poisson approximation for *P* (*M* = *m*) in equation Eq. (3) then yields

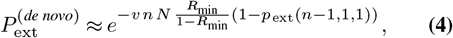

with which we recover the common approach to study evolutionary rescue (Orr and Unckless 2008; Martin *et al*. 2013; Uecker *et al*. 2014; Santer and Uecker 2020).

## Results

introresults To assess the probability of success of the different strategies, we begin by analyzing the initial sensitizing effects of the four pCRISPR variants on the various cell types present in the standing genetic varation (step 2 in our model with the initial cell-type distribution given through step 1). We then calculate the extinction probabilities under antibiotic treatment for single cells and afterwards for the entire population based on the birth-death process (step 3 in our model). Finally, we examine how the model parameters table 1 reference II(Table 1) affect these probabilities to determine which pCRISPR variant is most effective for specific conditions.

### A. Sensitizing effect

Prior to the introduction of pCRISPR (step 1), the overwhelming majority of cells carry only WT pAMR (referred to as WT cells in the following), a small fraction carries only pAMRmut (referred to as homozygous mutant cells), and some carry both, pAMRmut and WT pAMR copies (referred to as heterozygous cells). This distribution is shown as gray bars in Fig. 2 A.

Upon the introduction of pCRISPR (step 2), the WT cells become fully sensitized to antibiotics – all pAMR copies are cleaved or silenced immediately. However, this is not the case for heterozygous nor for homozygous mutant cells.

The distribution of pAMRmut copies in the population immediately after the introduction of pCRISPR and right before antibiotic treatment starts differs among the pCRISPR variants (Fig. 2 A):

*pCleaving:* pAMR copies in heterozygous cells are cleaved and replaced, either solely by pAMRmut copies (compatible pCRISPR) or both, pAMRmut and pCleaving copies (incompatible pCRISPR). Thus, for compatible pCleaving, more homozygous mutant cells will arise than for incompatible pCleaving, where pAMRmut and pCRISPR compete for replication.

*pSilencing:* immediately after the introduction the plasmid composition does not change (except for the addition of pSilencing).

At each cell division following pCRISPR introduction, the composition of pAMR, pAMRmut, and incompatible pCRISPR copies will change due to replication and segregation of the plasmids. If the population kept proliferating in the absence of antibiotics over multiple generations, segregation of cells that are heterozygous for incompatible plasmids into homozygous cells would occur. This emergence of homozygous cells occurs quite rapidly within a few generations (Ilhan *et al*. 2019). The proportion of the resulting homozygous cell types depends on the pCRISPR variant:

*For compatible pCleaving:* cells become AMR-plasmid-free or homozygous for pAMRmut.

*For incompatible pCleaving:* cells become homozygous for either pAMRmut or pCleaving.

*For compatible pSilencing:* cells become homozygous for pAMRmut or silenced pAMR, ignoring differential costs of plasmids and new mutations from pAMR to pAMRmut.

*For incompatible pSilencing:* cells become homozygous for one of pAMRmut, non-silenced pAMR, or pSilencing.

Ignoring any fitness differences and new mutations from pAMR to pAMRmut, the probabilities at which either homozygous cell type emerges in the population are given by the frequencies of the respective plasmid in *f* ^(CRISPR)^. The resulting cell type frequencies of pAMR, pAMRmut and pCRISPR are shown in Fig. 2 B.

In the following, we assume antibiotics are administered immediately after addition of pCRISPR without any delay, i.e. the cell type distribution is as in Fig. 2 A. If there was a short time gap between the introduction of pCRISPR and the administration of antibiotics, the distribution would start shifting towards the one shown in Fig. 2 B, before in the long run, the cost of pAMRmut would have an influence too.

### B. Extinction probabilities under treatment with antibiotics

The success probability of an antibiotic treatment that starts directly after the pCRISPR introduction depends on the pCRISPR variant. In the following, we first discuss single-cell extinction probabilities *p*_ext_ (ignoring new mutations) before we determine 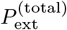, the extinction probability of the entire bacterial population for the different pCRISPR designs.

#### B.1. Single cell extinction probabilities

Since WT cells that contain only pAMR are the most prevalent at the start of the treatment, efficient sensitization of WT cells is a prerequisite for successful treatment. Only pCleaving and compatible pSilencing lead to high enough extinction probabilities of wild-type cells (Fig. 3 A). Since pCleaving eliminates all pAMR and compatible pSilencing permanently silences all of them, single-cell extinction probabilities are equal to one if we ignore that a new mutation could arise as we do in the calculation of *p*_ext_. (We will incorporate *de novo* mutations later.)

**Fig. 3.**
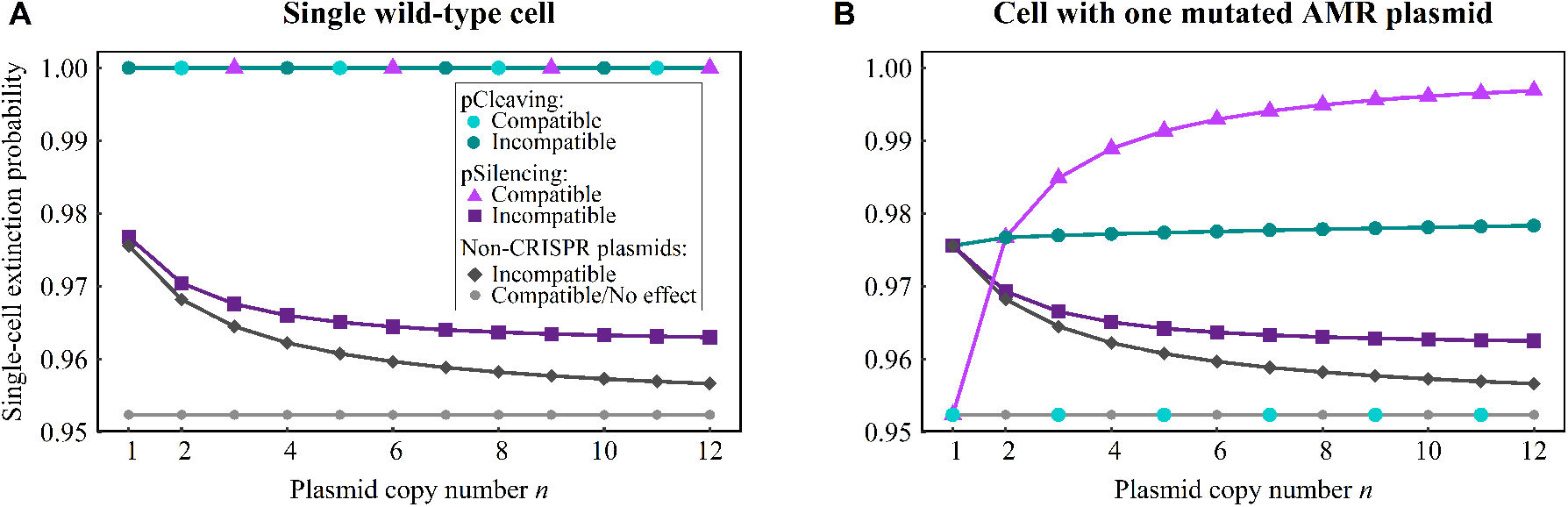
Extinction probabilities for a single cell lineage due to antibiotic treatment after pCRISPR introduction. The extinction probability for a lineage tracing back to a single wild-type cell (A) or a single cell carrying exactly one pAMRmut and *n* − 1 WT pAMR copies (B) prior to the CRISPR-mediated cleaving or silencing. For comparison, results for treatment with plasmids without a CRISPR-Cas system are also shown. Since a compatible non-CRISPR plasmid has no effect, this scenario is equivalent to the antibiotic treatment of a resistant population. Also note that, in the case of single wild-type cells (A), the extinction probability for pCleaving and also compatible pSilencing is precisely 1, as we exclude the tiny chance that a *de novo* mutation occurs for single cell lineage extinction probabilities and compute the probability of evolutionary rescue due to *de novo* mutations separately. For incompatible pSilencing and the non-CRISPR plasmids, a lineage extinction probability of at most 0.98 for WT cells is not high enough to treat a complete population of millions of cells. However, for the much lower number of mutated cells, the probabilities shown in (B) can be sufficient. The analysis uses a linear fitness function with parameters: *R* _max_ = 1.05, *R* _min_ = 0.95.

For incompatible pSilencing, plasmid competition, which is independent of CRISPR-Cas interference, can lead to the loss of pCRISPR and consequently to the termination of transcriptional silencing of the WT pAMR copies. Therefore, as already pointed out above, wild-type cells have a considerable chance of generating a long-term lineage of offspring without the need for *de novo* mutations. This renders incompatible pSilencing similarly inefficient as the introduction of an incompatible plasmid without a CRISPR-Cas system.

In contrast, a compatible/incompatible pCleaving and a compatible pSilencing, followed by antibiotic administration, would result in a successful treatment, if pAMR could not mutate to evade CRISPR-Cas interference. However, it is more realistic to assume that cells carrying mutated AMR plasmids pre-exist or in the case of compatible pSilencing arise *de novo* and may prevent the extinction of the population. For cells carrying at least one pAMRmut copy, whether they replicate or die crucially depends on their fitness after pCRISPR introduction, which in turn depends on the amount of functional AMR plasmids (Table 2) and the fitness function (dominant, linear, recessive). However, this fitness is not simply ‘passed on’ to the daughter cells but can change since plasmid replication and segregation change the relative abundance of functional AMR copies in the daughter cells with respect to the mother cell.

**Table 2.** Total number of functional AMR plasmids at the start of the antibiotic treatment (step 3) for a cell with *j* (≤ *n*) pAMRmut copies before pCRISPR introduction (step 2). For incompatible pCleaving, the cell contains *j* pAMRmut copies and one pCleaving copy after CRISPR-Cas interference. These are randomly replicated to *n* copies in total, and the value in the table is the expected amount of pAMRmut copies post replication. A similar argument holds for incompatible pSilencing. In the other two cases, the number is nonrandom. Furthermore, for compatible pCleaving, each originally heterozygous cell becomes immediately homozygous for pAMRmut. For both variants of pSilencing, the number of pAMR and pAMRmut copies does not change due to the introduction of pSilencing. 𝟙 is the indicator function. The replication model does not influence the numbers of functional AMR plasmids.

|  | pCleaving | pSilencing |
| --- | --- | --- |
| Incompatible | $nj/(j+1)$ | $nj/(n+1)$ |
| Compatible | $n \cdot \mathbb{1}_{j \neq 0}$ | $j$ |

The number of generations required for sensitized cells with at least one pAMRmut copy to restore sufficient antibiotic resistance depends besides on the fitness function also on the pCRISPR variant.

*Compatible pCleaving:* splittypestwo all pAMR copies within a heterozygous cell are cleaved and the number of pAMRmut copies increases to *n* (as stated in Table 2). Therefore, even a single pAMRmut copy fully prevents any sensitizing effect, making this design rather unsuitable to manage pre-existing variation.

*Incompatible pCleaving:* the immediate generation of homozygous pAMRmut cells out of heterozygous ones is prevented, and the extinction probability is thus higher than for compatible pCleaving.

*Incompatible and compatible pSilencing:* the immediate appearance of fully resistant cells from cells that were not homozygous for pAMRmut before is impossible as the silenced, i.e. transcriptional repressed, but replicating pAMR (and pSilencing) copies compete for replication with the pAMRmut copies.

We specifically analyze the long-term extinction probability *p*_ext_ of cells containing exactly one pAMRmut and *n* − 1 pAMR copies just prior to pCRISPR introduction (Fig. 3 B). These cells are particularly important, as they are for most copy numbers common in the standing genetic variation (compared to cells with more than one pAMRmut copy) and are the ones that arise from *de novo* mutations (at least in our approximation). Since we have already identified incompatible pSilencing as a poor strategy, and for pCleaving the compatible variant is less effective than the incompatible variant, we now focus on the two remaining strategies: compatible pSilencing and incompatible pCleaving.

n greater or equal 2 Whether compatible pSilencing or incompatible pCleaving is more successful in eradicating cells with one single pAMRmut copy depends on the copy number of pAMR (see Fig. 3 B and Fig. 4). In the case of incompatible pCleaving, two incompatible plasmids persist post-cleavage: one pCRISPR and one pAMRmut copy, which then replicate until *n* copies are present in the cell again. Consequently, the expected number of pAMRmut copies after plasmid reproduction is *n/*2. For compatible pSilencing, pAMR and pAMRmut copies are not cleaved. Thus, there is only one functional plasmid copy (the pAMRmut) after CRISPR-Cas introduction. In line with this consideration, incompatible pCleaving is superior for *n* = 1, while compatible pSilencing is superior for *n* ≥ 3 (Fig. 3 B).

**Fig. 4.**
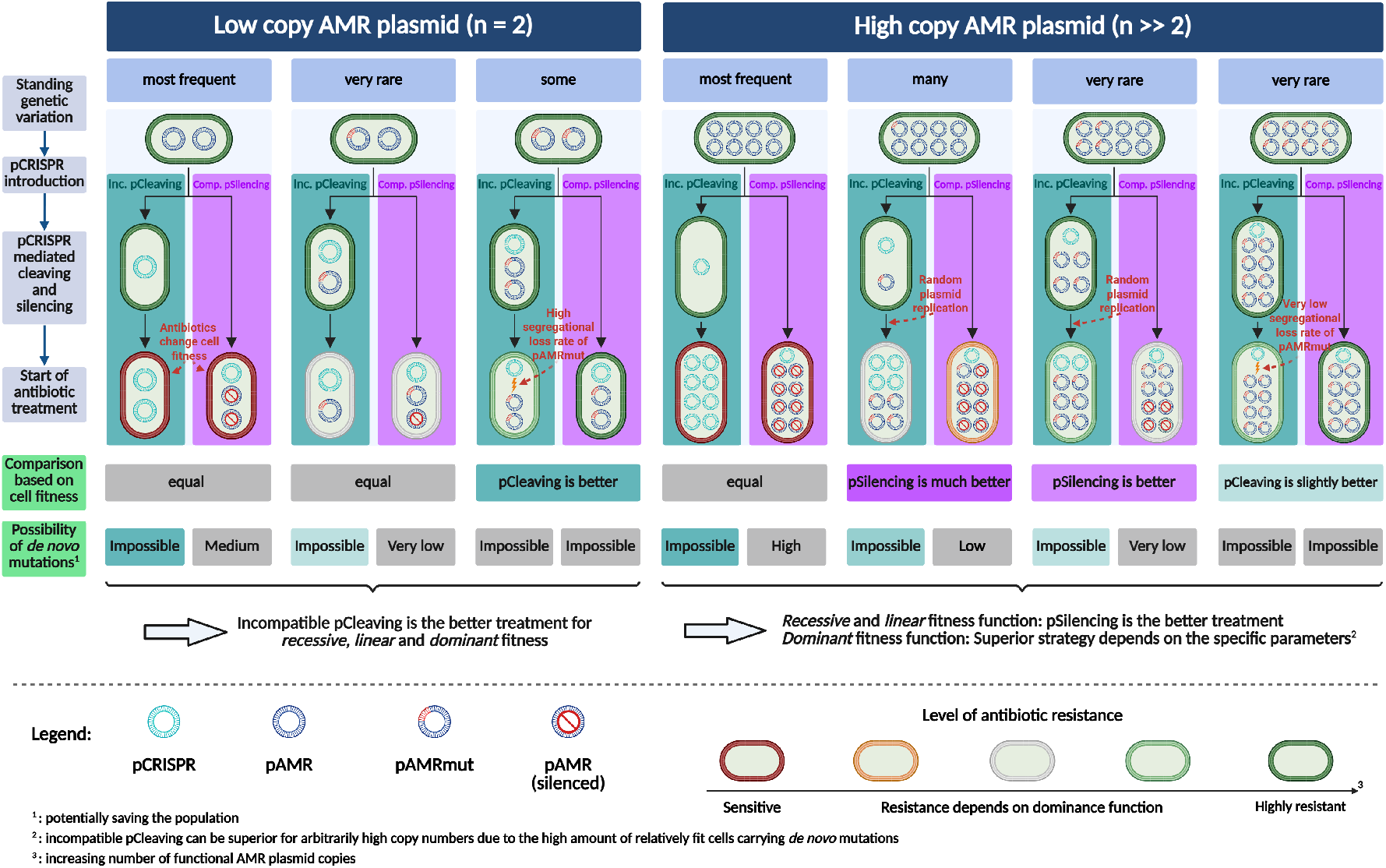
Theoretical comparison of the efficiencies of incompatible pCleaving and compatible pSilencing for a low-copy AMR plasmid (*n* = 2) and a high-copy AMR plasmid (*n* ≫ 2). The top row shows the frequency of exemplary cell types within the standing variation. The introduction of pCRISPR alters the plasmid composition within the cells, which depends on the CRISPR-Cas type (middle part of the figure). The fitnesses of the cells post CRISPR-Cas treatment provide information on the prospects of the two CRISPR-Cas treatments, which is summarized in the figure in the row ‘Comparison based on cell fitness’. The last row in the figure assesses the probability of *de novo* mutations from a silenced pAMR to a pAMRmut, as these mutations can prohibit the success of the treatment. When considering the combined effects of these factors on population survival, we find that incompatible pCleaving consistently outperforms compatible pSilencing for *n* = 2. For high copy numbers, the picture is more complicated. A quantitative analysis shows that for a *recessive* and *linear* fitness function, compatible pSilencing is more advantageous once a critical threshold plasmid copy number is exceeded. For a *dominant* fitness function, such a threshold may or may not exist. Created in BioRender. Baumdicker, F. (2025) https://BioRender.com/zvrvlwb

lastb1paragraph We next turn to heterozygous cells with an arbitrary number of pAMRmut copies. Right after the introduction of pCRISPR, compatible pSilencing leads to a lower average number of functional copies of the AMR gene in a cell than pCleaving as long as there are at least two pAMR copies in the cell (in other words, if the number *j* of pre-existing pAMRmut copies in the cell is less than *n* − 1, i.e. *j < n* − 1; see Table 2 and Fig. 4). When there are initially *n* − 1 pAMRmut copies in the cell, incompatible pCleaving and compatible pSilencing lead to the same average number of functional copies. shift for n greater 2 Consequently, for *n* = 2, heterozygous cells are equally well managed by both pCRISPR variants, but for larger copy numbers, nearly all heterozygous cells are better managed by compatible pSilencing. Mutant homozygous cells are always better managed by incompatible pCleaving than by compatible pSilencing due to competition between the resistance and the CRISPR-Cas plasmid in the former but not in the latter strategy.

### B.2 Extinction probabilities of the entire population

Our analysis of single-cell extinction probabilities revealed that CRISPR-Cas treatment using either incompatible pCleaving or compatible pSilencing are promising strategies and that the preferred approach depends on the cell type, i.e. whether the target plasmid is present in a homozygous or heterozygous state. To determine the total extinction probability of the bacterial population, we need to account for the distribution of cell types in the standing genetic variation and the risk of *de novo* mutations from pAMR to pAMRmut during population decline. From the previous section, it is already clear that incompatible pCleaving leads to a higher extinction probability than compatible pSilencing for *n* = 1 and *n* = 2. For higher copy numbers, it is less clear since compatible pSilencing is better at managing heterozygous cells (*j < n* − 1) or equally good (*j* = *n*− 1), but worse for homozygous mutant cells and moreover allows for rescue from *de novo* mutations.

The fraction of heterozygous or homozygous mutant cells in the standing genetic variation depends on the copy number (and also on the selection pressure prior to pCRISPR introduction, see section B.3). While the occurrence of mutated plasmid variants increases with the plasmid copy number, their within-cell fixation probability decreases. Therefore, higher copy numbers lead to a larger fraction of heterozygous cells but fewer homozygous cells (row ‘Standing genetic variation’ in Fig. 4 and Supplementary Fig. S5). Therefore, as long as we ignore *de novo* mutations of silenced pAMR copies, there always exists a threshold copy number above which compatible pSilencing is preferable over incompatible pCleaving (consistent with the ‘Comparison based on cell fitness’ in Fig. 4). However, *de novo* mutations can lower the effectiveness of compatible pSilencing (assessed in the last row of Fig. 4). Nevertheless, the example in Fig. 5 A shows that compatible pSilencing is still superior for copy numbers *n* ≥ 5 for these specific parameter choices. We find that the existence of a threshold, if we account for *de novo* mutations, depends on the fitness function (see Supplementary Note 3). In brief, for a linear and recessive fitness function, a threshold always exists. For dominant fitness functions, the existence of the threshold is possible, but not guaranteed; in the latter case, pCleaving is superior for all copy numbers *n*. Throughout the examples in this paper, a threshold always existed and was between a copy number of 4 and 10 but this will depend on the concrete parameters, as analysed in the next section.

**Fig. 5.**
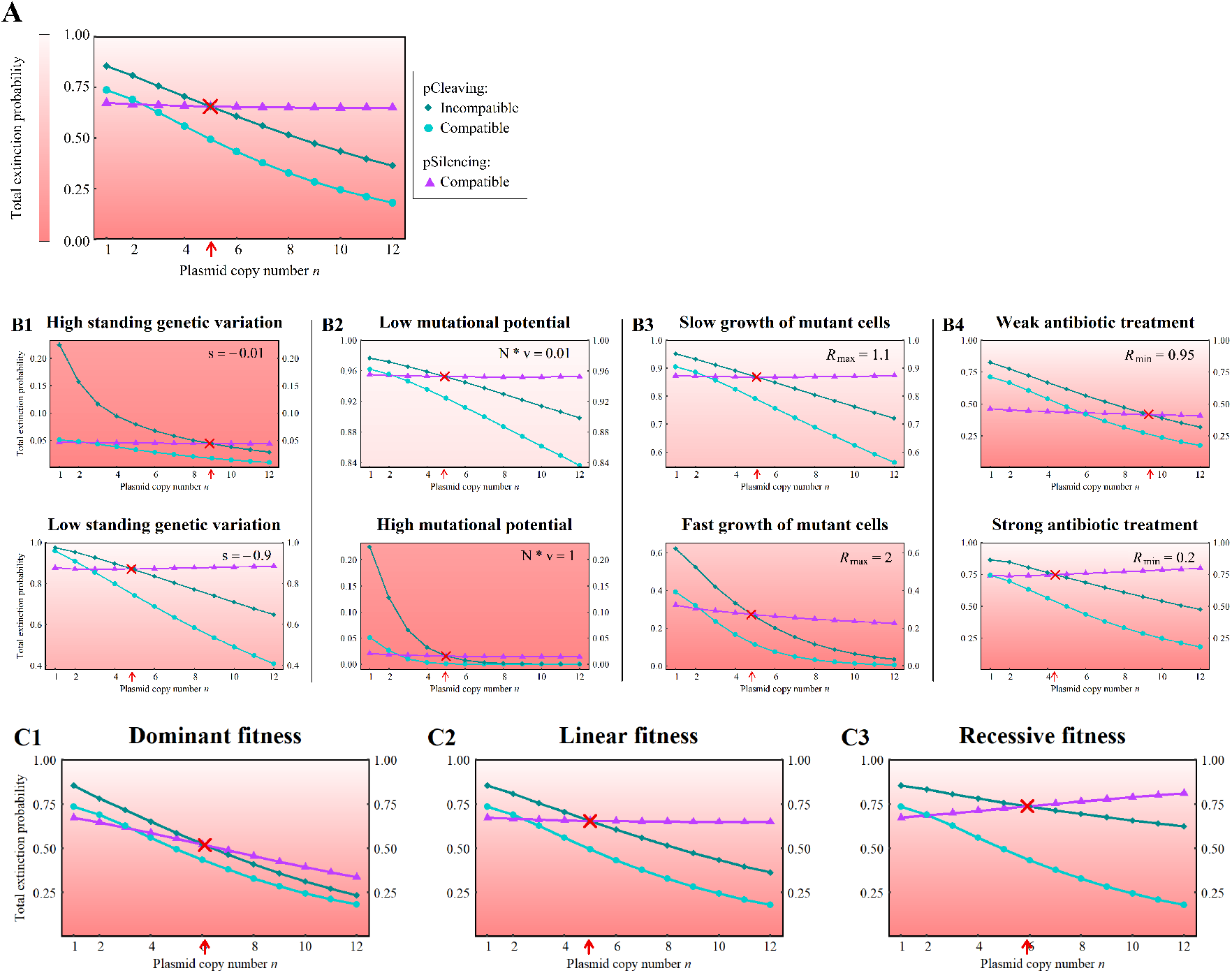
Total extinction probability in a pathogenic population. The focus is only on the following three primary cases: compatible and incompatible pCleaving and compatible pSilencing. Other cases from Fig. 3 are omitted due to their limited potential to eradicate the population. Panel (A) shows the influence of the copy number on the population extinction probability for all three cases. The parameters are *R* _max_ = 1.3, *R* _min_ = 0.8, *v* = 10^−8^, *s* = −0.1, *N* = 10^7^ assuming a linear fitness function. Panel (B) illustrates the effect when a specific parameter is adjusted to a higher or lower value as indicated in the top right-hand corner. All other parameters remain as in (A). In Panel (C), the fitness function is varied while all parameters are the same as in (A). The red arrow and cross indicate the point of intersection of the two curves representing compatible pSilencing and incompatible pCleaving. Below this intersection point, incompatible pCleaving is consistently superior, while above it, compatible pSilencing is more effective. Note the different ranges on the y-axis across panels. The gradient of the background color from white to light red visually represents the success of the treatment.

### B.3 The Effectiveness of pCleaving and pSilencing across various conditions

We next investigate the population extinction probability under various conditions and focus particularly on the threshold copy number above which compatible pSilencing is preferable over incompatible pCleaving. In Fig. 5 B, we alter a single model parameter with respect to the scenario shown in Fig. 5 A.

A pCRISPR targeting a more conserved DNA segment (i.e. strong selection *s* against pAMRmut prior to treatment) leads to fewer cells carrying pAMRmut copies in the standing genetic variation, thereby enhancing the probability of population extinction. As compatible pSilencing is most effective for cells with a low proportion of pAMRmut copies, this enhances the efficacy of compatible pSilencing compared to incompatible pCleaving, as the fraction of cells possessing a low count of pAMRmut copies increases. Consequently, the threshold occurs at a lower copy number than for weak negative selection against pAMRmut, as depicted in Fig. 5 B1.

A higher mutational potential, which is the product of the mutation rate and the initial population size, increases the occurrence of cells carrying pAMRmut copies, which leads to a lower chance of extinction of the bacterial population. Likewise, the extinction probability decreases with the growth rate of mutant cells. The threshold copy number is not strongly affected by either of these factors (Fig. 5 B2 and B3). This is because changes in the growth rate, the mutation rate, and the population size only alter the total number of cells with pAMRmut but not the composition of the standing genetic variation at the start of treatment. The larger number of new mutations with a higher mutational input is expected to decrease the relative effectiveness of compatible pSilencing compared to pCleaving, but this does not seem to affect the threshold much.

The intensity of the antibiotic treatment, i.e. the rate of decline of wild-type cells, has a particularly strong effect on the extinction probability when considering pSilencing. An increase in the intensity of antibiotic treatment leads to a faster population decline and consequently to a lower expected number of *de novo* mutations of silenced pAMR into functional pAMRmut copies. Moreover, with a linear fitness function as in Fig. 5 A and B, a strong treatment also entails a low fitness of heterozygous cells with few pAMRmut copies, which are more common for pSilencing than for pCleaving after pCRISPR has been introduced. Both factors do not have a strong effect for pCleaving, where *de novo* mutations are irrelevant and pAMRmut copies replicate to higher frequencies. Thus, the strength of the antibiotic treatment is an important parameter that determines whether pSilencing is better than pCleaving for a low or high copy number threshold, as can be seen in Fig. 5 B4.

To further investigate the effect of the fitness of heterozygous cells, we considered different fitness functions. If the maximum and minimum fitness of a cell remain the same, the fitness function (*dominant, linear* and *recessive*) only affects the fitness of cells where at least one but not all plasmid copies are functional pAMR or pAMRmut. Fig. 5 C shows that a dominant function results in the lowest extinction probability, while a recessive function leads to the highest. Compatible pSilencing is particularly interesting here, since with increasing copy number, both the number of *de novo* mutations and the fraction of cells in the SGV with only a few pAMRmut copies increases, while the fraction of homozygous mutant cells decreases (see Supplementary Fig. S5). The influence of the copy number on the overall extinction probability therefore depends on whether the contribution from many cells with a few pAMRmut copies (from *de novo* mutations or SGV) outweighs the contribution from few cells with solely pAMRmut copies. Consequently, the success rate of compatible pSilencing decreases monotonically with the copy number for dominant fitness, is almost constant for linear fitness, and increases monotonically for a recessive fitness function.

weirdly In contrast to compatible pSilencing, compatible pCleaving is completely unaffected by the fitness function as for the latter all pAMR are cleaved and potentially replaced by pAMRmut so that there are no heterozygous cells after CRISPR-Cas interference. For incompatible pCleaving, the extinction probability decreases with the plasmid copy number for all fitness functions, but extinction is most likely under a recessive fitness function. How the fitness function influences the population extinction probabilities is further illustrated in Supplementary Fig. S6.

## Discussion

In our study we have developed a mathematical model to explore strategies for combating bacteria resistant to conventional antibiotics using CRISPR-Cas technology. The model incorporates the dynamics of multicopy AMR plasmids and their interaction with different compatible and incompatible CRISPR-Cas-encoding plasmids, which either cleave or silence the targeted AMR genes. The success probability of the subsequent treatment is contingent on various factors, some of which are included in our analysis, while other potentially important factors are discussed in this section. In addition, we provide a model evaluation and validation broadly following the framework developed by Augusiak *et al*. (2014) in Supplementary Note 2.

Our results suggest that cleaving of the resistance gene by a CRISPR-Cas system encoded on a plasmid that is incompatible with the AMR plasmid is the most effective strategy if the pAMR has a low copy number. To manage pAMR of higher copy numbers, a CRISPR-Cas system for gene silencing encoded on a plasmid that is compatible with pAMR offers the superior solution under most circumstances. This result is supported by a mathematical argument in Supplementary Note 3 and robust to parameter variation (Fig. 5).

copynumber This implies that sequence specific CRISPR-Cas therapeutics require genetic information beyond the precise target sequence, which currently are not always available. First, the location of the resistance gene, i.e chromosome or plasmid, needs to be identified. This may be inferred from the resistance gene itself, for example by PCR testing (Anjum *et al*. 2017), as a given resistance gene is either typically associated with plasmids or the chromosome (Partridge *et al*. 2018). For a chromosomally encoded AMR gene, our results for *n* = 1, as well as the fact that cleaving the chromosome leads to cell death, both imply that pCleaving will generally be superior. Second, for plasmid-encoded AMR genes, the plasmid family (incompatibility group) needs to be identified in order to have a robust estimate for the copy number (Ramiro-Martínez *et al*. 2024). Nanopore sequencing has been shown to identify both, the resistance gene and the plasmid family within less than five hours from a positive blood culture (Taxt *et al*. 2020). Although not part of routine diagnostics, such analyses may be feasible in selected cases—for example, to sensitize a patient before surgery.

The incompatibility group is particularly important to evaluate the effectiveness of pSilencing. If pSilencing happens to be incompatible with pAMR, plasmid competition would frequently result in the loss of pSilencing, thereby stopping transcriptional repression and restoring AMR. It would therefore be necessary for incompatible pSilencing (and advantageous for incompatible pCleaving) to enhance its competitiveness and stability. This could be achieved by equipping pCRISPR, whether it acts by silencing or cleaving, with toxin-antitoxin (TA) systems (Kamruzzaman *et al*. 2017). toxin antitoxin These act as plasmid addiction systems by post-segregational killing of cells which have lost the TA system encoding plasmid (Hayes 2003; Jurėnas *et al*. 2022) and are encoded on many AMR plasmids to promote their stability (Yang and Walsh 2017). On the other hand, TA systems on the resistance encoding plasmids could enhance the effectiveness of cleaving pCRISPR, as bacteria from which the plasmid has been removed are readily killed by the toxin and cannot evolve resistance by other means such as mutations in the chromosome (a possibility that is not included in our model) (Citorik *et al*. 2014; Reuter *et al*. 2021).

The choice of the specific genetic target seems highly relevant. Targeting AMR genes directly is useful because escaping requires deleting/mutating this target sequence and arising pAMRmut copies are likely to have no or non-functional AMR genes. This contrasts with our model assumptions, where we ignore such nonfunctional mutations as they are not able to rescue the population from extinction. We thus included only mutations that evade CRISPR-Cas targeting while maintaining full functionality in our model. To further decrease the occurrence of pAMRmut with a functional AMR gene, several sites within the AMR gene, or other loci, can be targeted simultaneously. The effect of this strategy is covered by our modeling approach by setting a sufficiently low mutation probability. Our results show that lowering the mutational potential can dramatically increase the extinction probabilities (Fig. 5). Targeting other regions of the AMR plasmid in addition to or instead of the AMR gene is an interesting option. First, with the addition of more targets, mutating these simultaneously and thus escaping becomes increasingly unlikely. Second, depending on the chosen loci, targeting or escaping can result in decreased fitness and thus the loss of pAMRmut. A suitable target may be the plasmid replication machinery, which might hinder the proper replication of pAMRmut or slow down their replication. However, targeting the AMR plasmid backbone alone is likely not a promising strategy: AMR genes on plasmids are frequently encoded within mobile elements such as insertion sequences or transposition units, which can integrate into the chromosome or other plasmids (Partridge *et al*. 2018). Targeting the plasmid would select for AMR genes that have jumped to other replicons, whereas targeting the AMR gene would sensitize the cell independently of the gene’s location.

Mutations in the target sequence are not the only mechanism leading to failure of the treatment. Several experimental studies have demonstrated how the application of antibiotics following CRISPR-Cas interference selects for defective constructs (Bikard *et al*. 2014; Citorik *et al*. 2014; Reuter *et al*. 2021; Mamontov *et al*. 2022). This poses a real risk for treatment success, especially for applications with pSilencing, which needs to remain functional throughout antibiotic treatment. As a result, in reality, the probability of escape from pSilencing during treatment may be higher than assumed in our model. Using multiple copies of the *cas* genes (Li *et al*. 2023a) as well as reintroducing functional pCRISPR (Uribe *et al*. 2021) might reduce this risk. In Supplementary Fig. S7 we explore the scenario in which cells constantly acquire additional pCRISPR copies after the initial introduction. However, as we assume that initially every cell receives a pCRISPR, this has minimal impact on the results, even at high pCRISPR reinsertion rates.

For target interference by cleaving, Mamontov *et al*. (2022) discussed the existence of equilibria between CRISPR-Cas interference and pAMR replication rates. When both rates are equal, this can result in stable equilibria at numbers of copies below the actual copy number. However, experimental studies have shown that interference by cleaving was efficient even with plasmids present in up to 300 copies per cell (Citorik *et al*. 2014; Tagliaferri *et al*. 2020) and thus, in our model, we assume that CRISPR-Cas interference outweighs the pAMR replication rate.

While the assumption that the elimination of pAMR copies in cells which contain pCleaving is fast because of a high cleaving rate is reasonable, it is unlikely that the remaining pAMRmut and pCRISPR copies replicate at the same rate. A difference between the replication rates of pAMR and pCRISPR influences only the efficacy of the incompatible pCRISPR (Supplementary Fig. S8). If incompatible pCRISPR can replicate itself very quickly, it can be more effective than compatible pSilencing, even for higher copy numbers. Conversely, if pCRISPR replicates more slowly than the AMR plasmid, this reduces the extinction probability.

horizontal gene transfer Our analysis did not account for horizontal transfer of pCRISPR and for simplicity assumes that each cell receives a pCRISPR via an unspecified delivery mechanism. However, in reality, efficient *in situ* delivery of CRISPR-Cas tools is a main bottleneck for their application as novel antimicrobials. Recent efforts highlight a high potential of conjugative plasmids, delivering CRISPR-Cas tools to nearly 100% of the target bacterial population in a mouse colonisation model (Neil *et al*. 2020). HGT pCRISPR A conjugative pCRISPR could promote its own spread and pCRISPR could potentially be reintroduced to cells where (incompatible) pCRISPR has been lost. Using a conjugative incompatible pCleaving for delivery could be very efficient as pCleaving replaces all pAMR in recipient cells and thereby enhances its own spread. Using a transferable compatible pSilencing would also be beneficial. For pSilencing it is important to maintain the silencing effect and the reintroduction of a functional pSilencing could help to mitigate the effect of mutations that defect pSilencing.

HGT pAMR Horizontal transfer is not only an important consideration for the delivery plasmid but also for the AMR-encoding plasmids, which are frequently conjugative (i.e. self-transmissible) and can transfer to plasmid-free cells within the population. This suggests that cells that have lost the targeted plasmid may reacquire pAMR. However, it was shown that as long as these cells retain pCRISPR, they remain immune to reacquisition of pAMR (Rodrigues *et al*. 2019). In contrast, they are likely susceptible to pAMRmut, which can escape CRISPR-Cas interference, thereby lowering the extinction probability. The compatible pCleaving approach would be most affected by a conjugative pAMR/pAMRmut, as any wild-type cell acquiring a pAMRmut via conjugation would replicate these genes up to the original copy number *n*. This would further increase the number of pAMRmut and reinforce the transfer to additional cells. Incompatible pCleaving would be affected to a lesser extent. Compatible pSilencing would be least impacted, due to the lower number of pAMRmut copies in comparison to incompatible pCleaving and the presence of silenced pAMR in the same incompatibility group within the recipient cell.

limitations In this study, we focused on the population genetic aspects of the problem and kept the ecology as simple as possible. We assumed that the antibiotic pressure remains constant throughout the antibiotic treatment. If the antibiotic concentration intermittently dropped below the minimal inhibitory concentration, which *in vivo* could happen due to pharmacokinetic processes and/or skipped doses, the sensitive cell population could recover, and we would underestimate the risk for escape mutations for populations treated with pSilencing in which pAMR copies persist. spatial Moreover, our model assumes a well-mixed bacterial population and does not account for interactions between cells during treatment nor spatial structures like those in biofilms. Spatial structure may initially facilitate rapid plasmid spread, but for example the lack of regular mixing in biofilms likely prevents pCRISPR from reaching all cells within the population (Fox *et al*. 2008). Furthermore, the cell fitness can strongly depend on the composition of the neighboring cells. This would likely affect treatment success for different pCRISPR variants to different extents.

Although our model has been formulated in terms of plasmid-plasmid interactions, the results extend to chromosome-plasmid and plasmid-chromosome interactions and various gene delivery systems. A compatible CRISPR-Cas encoding plasmid that interacts with an AMR plasmid mirrors the dynamics of a chromosomal CRISPR-Cas system targeting an AMR plasmid. Vice versa, an AMR plasmid with a copy number of one mimics AMR genes that are encoded on the chromosome, as in our case, removal of the pAMR in a cell guarantees the extinction of its cell lineage, which is, at least in terms of the extinction probability, equivalent to killing the cell directly by cleaving the chromosome. For AMR genes on the chromosome, pCleaving is thus predicted to be more effective than pSilencing.

In conclusion, the innovative approach of sensitizing bacteria with CRISPR-Cas encoding plasmids and subsequently treating them with antibiotics is a promising technique in the fight against drug-resistant bacteria. This technique could redefine our ability to tackle microbial resistance and strengthen our arsenal in the global health battle. Our results apply to a variety of settings beyond direct treatment, such as reducing antibiotic resistance in aquacultures, improving community stability in bioreactors, or increasing the safety of fecal microbiome transplants (Valderrama *et al*. 2019; Li *et al*. 2023b). Our model suggests that treatment with a CRISPR-Cas-encoding plasmid incompatible with the target plasmid is only effective when using a DNA-cleaving interference mechanism. In contrast, a compatible pCRISPR is effective regardless of the underlying interference mechanism. However, transcriptional repression of the target gene is generally more effective for targeting plasmids with higher copy numbers, while cleaving is preferable for plasmids with lower copy numbers. These findings highlight that silencing mechanisms may provide a promising alternative to DNA-cleaving strategies, especially when targeting antimicrobial-resistant strains carrying high-copy-number plasmids.

## DATA AVAILABILITY

The authors state that all data necessary for confirming the conclusions presented in the article are represented fully within the article. The R-scripts for the calculations of the standing genetic variation, the extinction probabilities and more are available at https://github.com/fbaumdicker/SensitizingResistantBacteriaWithCRISPR.

## FUNDING

Johannes Kippnich is funded by the Deutsche Forschungsgemeinschaft (DFG, German Research Foundation) under the Priority Program - SPP 2141 - Project number Ba-5529/1-1 (to F.B.). Franz Baumdicker is funded by the Deutsche Forschungsgemeinschaft (DFG, German Research Foundation) under Germany’s Excellence Strategy – EXC number 2064/1 – Project number 390727645, and EXC 2124 – Project number 390838134. Fabienne Benz was supported by the SNSF (P500PB_210944).

## ACKNOWLEDGEMENTS

We thank Christin Nyhoegen for advice on the design of Fig. 1 and David Walker-Sünderhauf for insightful discussions.

## Notes

### Competing Interest Statement

The authors have declared no competing interest.

### Summary of Updates

Text updated to clarify model limitations. Figure 2 was added to better explain the plasmid copy number threshold.

## Bibliography

Adli M. 2018. The CRISPR tool kit for genome editing and beyond. Nature Communications. 9:1911.

Allen LJS. 2015. Stochastic Population and Epidemic Models: Persistence and Extinction. Springer International Publishing. Cham.

Anjum MF, Zankari E, Hasman H. 2017. Molecular Methods for Detection of Antimicrobial Resistance. Microbiology Spectrum. 5:10.1128/microbiolspec.arba–0011–2017.

Antimicrobial Resistance Collaborators. 2022. Global burden of bacterial antimicrobial resistance in 2019: a systematic analysis. Lancet. 399:629–655.

Augusiak J, Van den Brink PJ, Grimm V. 2014. Merging validation and evaluation of ecological models to ‘evaludation’: A review of terminology and a practical approach. Ecological Modelling. 280:117–128.

Benz F, Beamud B, Laurenceau R, Maire A, Duportet X, Decrulle A, Bikard D. 2025. CRISPR–Cas therapies targeting bacteria. Nature Reviews Bioengineering. pp. 1–18.

Bikard D, Euler CW, Jiang W, Nussenzweig PM, Goldberg GW, Duportet X, Fischetti VA, Marraffini LA. 2014. Exploiting CRISPR-Cas nucleases to produce sequence-specific antimicrobials. Nature Biotechnology. 32:1146–1150.

Bouet JY, Nordström K, Lane D. 2007. Plasmid partition and incompatibility – the focus shifts. Molecular Microbiology. 65:1405–1414.

Brödel AK, Charpenay L, Galtier M, Fuche FJ, Terrasse R, Poquet C, Arraou M, Prevot G, Spadoni D, Hessel EM et al. 2022. In situ targeted mutagenesis of gut bacteria. biorxiv: 2022.09.30.509847.

Buckner MMC, Ciusa ML, Piddock LJV. 2018. Strategies to combat antimicrobial resistance: anti-plasmid and plasmid curing. FEMS Microbiology Reviews. 42:781–804.

Carattoli A. 2009. Resistance plasmid families in Enterobacteriaceae. Antimicrobial Agents and Chemotherapy. 53:2227–2238.

Castañeda-Barba S, Top EM, Stalder T. 2024. Plasmids, a molecular cornerstone of antimicrobial resistance in the One Health era. Nature Reviews Microbiology. 22:18–32.

Citorik RJ, Mimee M, Lu TK. 2014. Sequence-specific antimicrobials using efficiently delivered RNA-guided nucleases. Nature Biotechnology. 32:1141–1145.

Dewan I, Uecker H. 2023. A mathematician’s guide to plasmids: an introduction to plasmid biology for modellers: This article is part of the Microbial Evolution collection. Microbiology. 169.

Dewan I, Uecker H. 2024. Evolutionary rescue of bacterial populations by heterozygosity on multicopy plasmids. biorxiv: 2024.05.29.596466.

Ebersbach G, Gerdes K. 2005. Plasmid segregation mechanisms. Annual Review of Genetics. 39:453–479.

Fox RE, Zhong X, Krone SM, Top EM. 2008. Spatial structure and nutrients promote invasion of IncP-1 plasmids in bacterial populations. The ISME Journal. 2:1024–1039.

Friehs K. 2004. Plasmid Copy Number and Plasmid Stability, In: Scheper T, editor, New Trends and Developments in Biochemical Engineering, Springer. Berlin, Heidelberg. Advances in Biochemical Engineering. pp. 47–82.

Garoña A, Santer M, Hülter NF, Uecker H, Dagan T. 2023. Segregational drift hinders the evolution of antibiotic resistance on polyploid replicons. Pages: 2023.02.01.526651 Section: New Results.

Gencay YE, Jasinskytė D, Robert C, Semsey S, Martínez V, Petersen AO, Brunner K, de Santiago Torio A, Salazar A, Turcu IC et al. 2024. Engineered phage with antibacterial CRISPR–Cas selectively reduce E. coli burden in mice. Nature Biotechnology. 42:265–274.

Grant AJ, Restif O, McKinley TJ, Sheppard M, Maskell DJ, Mastroeni P. 2008. Modelling within-Host Spatiotemporal Dynamics of Invasive Bacterial Disease. PLOS Biology. 6:e74. Publisher: Public Library of Science.

Grimm V, Berger U, Bastiansen F, Eliassen S, Ginot V, Giske J, Goss-Custard J, Grand T, Heinz SK, Huse G et al. 2006. A standard protocol for describing individual-based and agent-based models. Ecological Modelling. 198:115–126.

Grimm V, Railsback SF, Vincenot CE, Berger U, Gallagher C, DeAngelis DL, Edmonds B, Ge J, Giske J, Groeneveld J et al. 2020. The ODD Protocol for Describing Agent-Based and Other Simulation Models: A Second Update to Improve Clarity, Replication, and Structural Realism. Journal of Artificial Societies and Social Simulation. 23:7.

Guzmán-Herrador DL, Fernández-Gómez A, Depardieu F, Bikard D, Llosa M. 2024. Delivery of functional Cas:DNA nucleoprotein complexes into recipient bacteria through a type IV secretion system. Proceedings of the National Academy of Sciences. 121:e2408509121.

Hamilton TA, Pellegrino GM, Therrien JA, Ham DT, Bartlett PC, Karas BJ, Gloor GB, Edgell DR. 2019. Efficient interspecies conjugative transfer of a CRISPR nuclease for targeted bacterial killing. Nature Communications. 10:4544. Publisher: Nature Publishing Group.

Hayes F. 2003. Toxins-Antitoxins: Plasmid Maintenance, Programmed Cell Death, and Cell Cycle Arrest. Science. 301:1496–1499.

Hernandez-Beltran JCR, Rodríguez-Beltrán J, Aguilar-Luviano OB, Velez-Santiago J, Mondragón-Palomino O, MacLean RC, Fuentes-Hernández A, San Millán A, Peña-Miller R. 2024. Plasmid-mediated phenotypic noise leads to transient antibiotic resistance in bacteria. Nature Communications. 15:1–13.

Ilhan J, Kupczok A, Woehle C, Wein T, Hülter NF, Rosenstiel P, Landan G, Mizrahi I, Dagan T. 2019. Segregational Drift and the Interplay between Plasmid Copy Number and Evolvability. Molecular Biology and Evolution. 36:472–486.

Jin WB, Li TT, Huo D, Qu S, Li XV, Arifuzzaman M, Lima SF, Shi HQ, Wang A, Putzel GG et al. 2022. Genetic manipulation of gut microbes enables single-gene interrogation in a complex microbiome. Cell. 185:547–562.e22.

Jurėnas D, Fraikin N, Goormaghtigh F, Van Melderen L. 2022. Biology and evolution of bacterial toxin–antitoxin systems. Nature Reviews Microbiology. 20:335–350. Publisher: Nature Publishing Group.

Kaiser P, Slack E, Grant AJ, Hardt WD, Regoes RR. 2013. Lymph Node Colonization Dynamics after Oral Salmonella Typhimurium Infection in Mice. PLOS Pathogens. 9:e1003532. Publisher: Public Library of Science.

Kamruzzaman M, Shoma S, Thomas CM, Partridge SR, Iredell JR. 2017. Plasmid interference for curing antibiotic resistance plasmids in vivo. PLOS ONE. 12:e0172913.

Lam KN, Spanogiannopoulos P, Soto-Perez P, Alexander M, Nalley MJ, Bisanz JE, Nayak RR, Weakley AM, Yu FB, Turnbaugh PJ. 2021. Phage-delivered CRISPR-Cas9 for strain-specific depletion and genomic deletions in the gut microbiome. Cell Reports. 37.

Li Q, Sun M, Lv L, Zuo Y, Zhang S, Zhang Y, Yang S. 2023a. Improving the Editing Efficiency of CRISPR-Cas9 by Reducing the Generation of Escapers Based on the Surviving Mechanism. ACS Synthetic Biology. 12:672–680. Publisher: American Chemical Society.

Li X, Bao N, Yan Z, Yuan XZ, Wang SG, Xia PF. 2023b. Degradation of Antibiotic Resistance Genes by VADER with CRISPR-Cas Immunity. Applied and Environmental Microbiology. 89:e00053–23. Publisher: American Society for Microbiology.

Luepke KH, Suda KJ, Boucher H, Russo RL, Bonney MW, Hunt TD, Mohr III JF. 2017. Past, Present, and Future of Antibacterial Economics: Increasing Bacterial Resistance, Limited Antibiotic Pipeline, and Societal Implications. Pharmacotherapy: The Journal of Human Pharmacology and Drug Therapy. 37:71–84.

Mamontov V, Martynov A, Morozova N, Bukatin A, Staroverov DB, Lukyanov KA, Ispolatov Y, Semenova E, Severinov K. 2022. Persistence of plasmids targeted by CRISPR interference in bacterial populations. Proceedings of the National Academy of Sciences. 119:e2114905119.

Martin G, Aguilée R, Ramsayer J, Kaltz O, Ronce O. 2013. The probability of evolutionary rescue: towards a quantitative comparison between theory and evolution experiments. Philosophical Transactions of the Royal Society B: Biological Sciences. 368:20120088.

Mayorga-Ramos A, Zúñiga-Miranda J, Carrera-Pacheco SE, Barba-Ostria C, Guamán LP. 2023. CRISPR-Cas-Based Antimicrobials: Design, Challenges, and Bacterial Mechanisms of Resistance. ACS Infectious Diseases. 9:1283–1302.

Naghavi M, Vollset SE, Ikuta KS, Swetschinski LR, Gray AP, Wool EE, Aguilar GR, Mestrovic T, Smith G, Han C et al. 2024. Global burden of bacterial antimicrobial resistance 1990–2021: a systematic analysis with forecasts to 2050. The Lancet. 404:1199–1226.

Neil K, Allard N, Grenier F, Burrus V, Rodrigue S. 2020. Highly efficient gene transfer in the mouse gut microbiota is enabled by the Incl2 conjugative plasmid TP114. Communications Biology. 3:1–9.

Neil K, Allard N, Roy P, Grenier F, Menendez A, Burrus V, Rodrigue S. 2021. High-efficiency delivery of CRISPR-Cas9 by engineered probiotics enables precise microbiome editing. Molecular Systems Biology. 17:e10335.

Nicoloff H, Hjort K, Andersson DI, Wang H. 2024. Three concurrent mechanisms generate gene copy number variation and transient antibiotic heteroresistance. Nature Communications. 15:3981. Publisher: Nature Publishing Group.

Nordström K. 2006. Plasmid R1 - Replication and its control. Plasmid. 55:1–26.

Novick RP, Hoppensteadt FC. 1978. On plasmid incompatibility. Plasmid. 1:421–434.

Nussenzweig PM, Marraffini LA. 2020. Molecular Mechanisms of CRISPR-Cas Immunity in Bacteria. Annual Review of Genetics. 54:93–120.

Orr HA, Unckless RL. 2008. Population extinction and the genetics of adaptation. The American Naturalist. 172:160–169.

Park JY, Moon BY, Park JW, Thornton JA, Park YH, Seo KS. 2017. Genetic engineering of a temperate phage-based delivery system for CRISPR/Cas9 antimicrobials against Staphylococcus aureus. Scientific Reports. 7:44929.

Partridge SR, Kwong SM, Firth N, Jensen SO. 2018. Mobile Genetic Elements Associated with Antimicrobial Resistance. Clinical Microbiology Reviews. 31:10.1128/cmr.00088–17.

Pólya G. 1930. Sur quelques points de la théorie des probabilités. Annales de l’institut Henri Poincaré. 1:117–161.

Ramiro-Martínez P, Quinto Id, Gama JA, Rodríguez-Beltrán J. 2024. Universal rules govern plasmid copy number. Pages: 2024.10.04.616648 Section: New Results.

Reuter A, Hilpert C, Dedieu-Berne A, Lematre S, Gueguen E, Launay G, Bigot S, Lesterlin C. 2021. Targeted-antibacterial-plasmids (TAPs) combining conjugation and CRISPR/Cas systems achieve strain-specific antibacterial activity. Nucleic Acids Research. 49:3584–3598.

Rodrigues M, McBride SW, Hullahalli K, Palmer KL, Duerkop BA. 2019. Conjugative delivery of CRISPR-Cas9 for the selective depletion of antibiotic-resistant enterococci. bioRxiv. pp. 1–14.

Rodriguez-Beltran J, Sorum V, Toll-Riera M, de la Vega C, Pena-Miller R, San Millan A. 2020. Genetic dominance governs the evolution and spread of mobile genetic elements in bacteria. Proceedings of the National Academy of Sciences. 117:15755–15762.

Ronda C, Chen SP, Cabral V, Yaung SJ, Wang HH. 2019. Metagenomic engineering of the mammalian gut microbiome in situ. Nature Methods. 16:167–170.

Rozwandowicz M, Brouwer MSM, Fischer J, Wagenaar JA, Gonzalez-Zorn B, Guerra B, Mevius DJ, Hordijk J. 2018. Plasmids carrying antimicrobial resistance genes in Enterobacteriaceae. Journal of Antimicrobial Chemotherapy. 73:1121–1137.

San Millan A, Escudero JA, Gifford DR, Mazel D, MacLean RC. 2016. Multicopy plasmids potentiate the evolution of antibiotic resistance in bacteria. Nature Ecology & Evolution. 1:10.

Santer M, Kupczok A, Dagan T, Uecker H. 2022. Fixation dynamics of beneficial alleles in prokaryotic polyploid chromosomes and plasmids. Genetics. 222:iyac121.

Santer M, Uecker H. 2020. Evolutionary Rescue and Drug Resistance on Multicopy Plasmids. Genetics. 215:847–868.

Santos-Lopez A, Bernabe-Balas C, Ares-Arroyo M, Ortega-Huedo R, Hoefer A, San Millan A, Gonzalez-Zorn B. 2017. A Naturally Occurring Single Nucleotide Polymorphism in a Multicopy Plasmid Produces a Reversible Increase in Antibiotic Resistance. Antimicrobial Agents and Chemotherapy. 61:10.1128/aac.01735–16.

Schrader SM, Vaubourgeix J, Nathan C. 2020. Biology of antimicrobial resistance and approaches to combat it. Science translational medicine. 12:eaaz6992.

Selle K, Barrangou R. 2015. Harnessing CRISPR–Cas systems for bacterial genome editing. Trends in Microbiology. 23:225–232.

Shao B, Rammohan J, Anderson DA, Alperovich N, Ross D, Voigt CA. 2021. Single-cell measurement of plasmid copy number and promoter activity. Nature Communications. 12:1475.

Stougaard P, Molin S, Nordström K. 1979. Plasmid R1 in Salmonella typhimurium: Molecular instability and gene dosage effects. Plasmid. 2:589–597.

Tagliaferri TL, Guimarães NR, Pereira MdPM, Vilela LFF, Horz HP, dos Santos SG, Mendes TAdO. 2020. Exploring the Potential of CRISPR-Cas9 Under Challenging Conditions: Facing High-Copy Plasmids and Counteracting Beta-Lactam Resistance in Clinical Strains of Enterobacteriaceae. Frontiers in Microbiology. 11:1–11.

Taxt AM, Avershina E, Frye SA, Naseer U, Ahmad R. 2020. Rapid identification of pathogens, antibiotic resistance genes and plasmids in blood cultures by nanopore sequencing. Scientific Reports. 10:7622. Publisher: Nature Publishing Group.

Turkeltaub L, Kashat L, Assous MV, Adler K, Bar-Meir M. 2024. Estimating bacterial load in S. aureus and E. coli bacteremia using bacterial growth graph from the continuous monitoring blood culture system. European Journal of Clinical Microbiology & Infectious Diseases. 43:1931–1938.

Uecker H, Otto SP, Hermisson J. 2014. Evolutionary Rescue in Structured Populations. The American Naturalist. 183:E17–E35.

Uribe RV, Rathmer C, Jahn LJ, Ellabaan MMH, Li SS, Sommer MOA. 2021. Bacterial resistance to CRISPR-Cas antimicrobials. Scientific Reports. 11:17267.

Valderrama JA, Kulkarni SS, Nizet V, Bier E. 2019. A bacterial gene-drive system efficiently edits and inactivates a high copy number antibiotic resistance locus. Nature Communications. 10:5726. Publisher: Nature Publishing Group.

von Wintersdorff CJH, Penders J, van Niekerk JM, Mills ND, Majumder S, van Alphen LB, Savelkoul PHM, Wolffs PFG. 2016. Dissemination of Antimicrobial Resistance in Microbial Ecosystems through Horizontal Gene Transfer. Frontiers in Microbiology. 7:173.

Walker-Sünderhauf D, Klümper U, Pursey E, Westra ER, Gaze WH, van Houte S. 2023. Removal of AMR plasmids using a mobile, broad host-range CRISPR-Cas9 delivery tool. Microbiology. 169:001334.

Wang P, He D, Li B, Guo Y, Wang W, Luo X, Zhao X, Wang X. 2019. Eliminating mcr-1-harbouring plasmids in clinical isolates using the CRISPR/Cas9 system. Journal of Antimicrobial Chemotherapy. 74:2559–2565.

Yang QE, Walsh TR. 2017. Toxin–antitoxin systems and their role in disseminating and maintaining antimicrobial resistance. FEMS Microbiology Reviews. 41:343–353.

Yao S, Wei D, Tang N, Song Y, Wang C, Feng J, Zhang G. 2022. Efficient Suppression of Natural Plasmid-Borne Gene Expression in Carbapenem-Resistant Klebsiella pneumoniae Using a Compact CRISPR Interference System. Antimicrobial Agents and Chemotherapy. 66:e00890–22.

Yosef I, Manor M, Kiro R, Qimron U. 2015. Temperate and lytic bacteriophages programmed to sensitize and kill antibiotic-resistant bacteria. Proceedings of the National Academy of Sciences. 112:7267–7272.

Zhou Y, Yang Y, Li X, Tian D, Ai W, Wang W, Wang B, Kreiswirth BN, Yu F, Chen L et al. 2023. Exploiting a conjugative endogenous CRISPR-Cas3 system to tackle multidrug-resistant Klebsiella pneumoniae. eBioMedicine. 88:104445.

