## Supplementary Material for "Effectiveness of CRISPR-Cas in Sensitizing Bacterial Populations with Plasmid-Encoded Antimicrobial Resistance"

### Supplementary Note 1: ODD Protocol

Here we provide a description of our model that follows the ODD (Overview, Design concepts, Details) protocol for describing individual- and agent-based models (Grimm *et al.* 2006), as updated by Grimm *et al.* (2020) to provide an additional clear and structured overview of the model.

**Purpose and patterns.** This model aims to evaluate the effectiveness of CRISPR-Cas in sensitizing bacterial populations and to assess the probability of successful eradication through subsequent antibiotic treatment. It functions as a minimal conceptual framework, aimed at exploring the dynamics between plasmid-mediated antibiotic resistance and CRISPR-Cas interventions under generalized conditions – rather than being tailored for specific clinical scenarios such as individual patient treatment. The model explicitly accounts for different target interference mechanisms, the copy number of the resistance-encoding plasmid, and its compatibility with the CRISPR-Cas encoding plasmid, ensuring a comprehensive representation of the interplay between genetic resistance and CRISPR-based interventions.

We define several patterns that the model is designed to reflect:

(i) deleterious CRISPR-Cas evading plasmid mutations exist at low frequency due to mutation-selection dynamics (Santer and Uecker 2020),

(ii) plasmid replication and segregation follow stochastic rules (Novick and Hoppensteadt 1978),

(iii) plasmid incompatibility leads to competitive plasmid loss (Novick and Hoppensteadt 1978; Ebersbach and Gerdes 2005),

(iv) antibiotic treatment effectively eliminates cells without antimicrobial resistance (Schrader *et al.* 2020), and

(v) the level of antibiotic resistance of each cell depends on the plasmid composition in the cell (Rodriguez-Beltran *et al.* 2020).

**Entities, State Variables and Scales.** The model represents a bacterial population where each individual cell is characterized by its plasmid composition. Bacterial cells serve as the primary entities in the model, and their properties are governed by three plasmid types and their interactions:

- pAMR (Wild-type AMR plasmid): Provides the host cell with AMR.
- pAMRmut (Mutated AMR plasmid): A variant of pAMR that evades CRISPR-Cas targeting due to a mutation in the target gene. In a worst-case scenario, it confers the same level of antibiotic resistance as pAMR.
- pCRISPR (CRISPR-Cas-encoding plasmid): Enables CRISPR-Cas interference against pAMR but does not affect pAMRmut. Two distinct interference mechanisms are considered:
  - pCleaving: Induces double-strand breaks at the target DNA site.
  - pSilencing: Inhibits gene expression through transcriptional repression.

The plasmid composition inside each cell determines its state variables: the birth and death rates.

*Temporal scale:* The model operates in continuous time, where cellular processes such as division and death occur as stochastic events with exponentially distributed waiting times.

*Spatial scale:* The model assumes no spatial structure, and cells evolve independently without direct interactions.

**Process overview.** The model consists of three steps:

**Step 1 ($t < 0$): Standing genetic variation prior to any treatment:**
In this initial stage before the introduction of pCRISPR and antibiotic treatment, cells can harbor pAMR and/or pAMRmut. We assume that the CRISPR-Cas evading mutation introduces a small cost $s$ and that the population is in mutation-selection balance. The population size is constant. This is achieved by removing a random cell when another cell replicates.

**Step 2 ($t = 0$): Sensitizing via pCRISPR introduction:**
At the start of this treatment step, each cell acquires exactly one pCRISPR in addition to the existing pAMR and pAMRmut. The effect of pCRISPR on pAMR is assumed to be instantaneous, i.e. the WT pAMR copies are immediately cleaved or silenced.

**Step 3 ($t > 0$): Treating the population with antibiotics:**

The antibiotic treatment starts as soon as each cell has received pCRISPR. Cells without pAMRmut are sensitive to antibiotic treatment, as all pAMR are cleaved or silenced. If pSilencing is lost through plasmid segregation at cell division, all pAMR are immediately providing resistance for its host again. The survival probability of cell lineages that originate from cells with pre-existing or *de novo* acquired pAMRmut copies depends on demographic stochasticity, the number of pAMRmut copies, and the mode of action and compatibility type of pCRISPR.

**Design concepts.** *Basic principles:* The model addresses a fundamental problem in evolutionary biology: whether a population will go extinct or adapts. Its survival depends on the emergence of a previously costly mutation, which becomes essential once pCRISPR begins targeting the wild-type resistance. The goal is to estimate the extinction probability of the population, based on key parameters such as cell fitness and the specific pCRISPR variant used.

*Emergence:* In this model, the survival or extinction of the bacterial population results from the interplay between individual bacterial fitness, the cost and benefit of mutations, and the selective pressure imposed by pCRISPR. The emergence and spread of a key mutation necessary for survival is a direct result of adaptive evolutionary dynamics rather than an imposed rule. The probability of this mutation spreading depends on agent-level factors such as cell fitness, mutation rates, and plasmid segregation, as well as environmental conditions like antibiotic pressure. In contrast, the introduction of pCRISPR itself and its targeting efficiency are imposed by model parameters.

*Adaptation:* Adaptation occurs only through the evolutionary selection of mutations that influence bacterial survival under pCRISPR pressure. The model does not include any adaptive behavior in a sense where the agents, i.e. the cells, would have to make decisions.

*Interaction:* In our main model during treatment, individual cells do not interact with each other. This simplification was made to keep the model as straightforward and analytically tractable as possible while still allowing for meaningful conclusions. In particular, there is no competition for resources during treatment, meaning that the fitness of cells is not influenced by the population density. Prior to treatment, the population size is kept constant, which means that cells compete for resources. We discuss the potential impact of incorporating horizontal gene transfer in the discussion section. There, we discuss the consequences of allowing pAMR, pAMRmut, and/or pCRISPR to be copied and transferred horizontally between cells. Spatial structure is not taken into account in these considerations, so horizontal transfer occurs randomly between cells rather than being dependent on proximity.

*Stochasticity:* Stochasticity plays a crucial role in this model: Mutations from pAMR to pAMRmut can occur during plasmid replication, with a fixed probability $v$ at each plasmid replication. Consequently, the number of new mutations per division follows a binomial distribution. Plasmid copies are randomly picked for replication in the main model variant and are randomly partitioned between daughter cells at cell division. This introduces stochasticity in the plasmid dynamics and leads to distinct behaviors for high and low copy plasmids in terms of emergence of homozygous cells. The cell dynamics are subject to stochasticity as well and are governed by cell-type-specific birth and death rates, which define the exponentially distributed waiting times for these events. These stochastic elements enable the model to capture the inherent randomness of mutation-driven adaptation.

*Observation:* The model predicts the extinction probability (and thus the survival probability) of single cells and entire bacterial populations given all relevant parameters. To analyze these probabilities, different parameter sets were investigated, and the resulting data were used to generate Figures 2, 3, and 5. The code for these analyses is available at https://github.com/fbaumdicker/SensitizingResistantBacteriaWithCRISPR.

**Initialization.** At the start of step 1, all cells initially contain exactly $n$ copies of pAMR, corresponding to the predefined copy number. As cell divisions occur, pAMRmut variants emerge due to mutations. By the beginning of step 2, the population is assumed to have reached mutation-selection balance as a result of the processes in step 1.

**Input data.** The input data are the global parameters for each simulation, i.e. the fixed copy number $n$, the number of cells $N$, the cost of the mutation before pCRISPR introduction $s$ and the maximum and minimum fitness/birth-death ratio $R_{\text{max}}$ and $R_{\text{min}}$ as well as the fitness function, i.e. *dominant*, *linear* or *recessive*.

**Submodels.**

***Copy number and plasmid incompatibility.*** Throughout every phase of the model – and regardless of the pCRISPR compatibility scenario – we assume that each cell contains exactly $n$ copies of plasmids that are incompatible. We examine two distinct cases for pCRISPR: one in which pCRISPR is incompatible with pAMR/pAMRmut and another where it is compatible with them. pAMR and pAMRmut are assumed to be intrinsically incompatible due to their close relatedness. By denoting the number of pAMR copies in a cell as $\#$pAMR (and using similar notation for the other plasmids), the following conditions apply depending on the compatibility between the CRISPR-Cas and resistance plasmids:

**Incompatible pCRISPR**
$\#\text{pAMR} + \#\text{pAMRmut} + \#\text{pCRISPR} = n$ and

**Compatible pCRISPR**
$\#\text{pAMR} + \#\text{pAMRmut} = n.$

Note that in the compatible pCRISPR scenario, the number of pCRISPR copies does not affect the model dynamics as long as the plasmid is stably maintained.

***Plasmid replication and segregation during cell division.*** During each cell division, all plasmids that are incompatible with pAMR, including the pAMR copies themselves, replicate until the total count reaches $2n$. We consider two different plasmid replication mechanisms: one where the system retains memory of which plasmids have already been copied, and another where replication follows a memoryless process, as described in Novick and Hoppensteadt (1978) and Nordström (2006):

***Regular* replication:** Each plasmid is copied exactly once.

***Random* replication:** One incompatible plasmid at a time is randomly selected for replication and added to the pool until the copy number is reached.

Once replication is complete, the cell divides, ensuring that each daughter cell inherits precisely $n$ plasmid copies. The allocation of plasmids between the two daughter cells follows a hypergeometric distribution.

***Step 1: Moran model before pCRISPR introduction.*** We consider a constant population of $N$ bacterial cells, each harboring exactly $n$ copies of resistance plasmids. Initially, all plasmids are of type pAMR, so there are no pAMRmut or pCRISPR copies (i.e., # pAMRmut = # pCRISPR = 0). Each cell divides at a certain rate depending on its type. When a cell divides, the mother cell produces two daughter cells, replacing both itself and a randomly chosen cell to maintain a constant population size. We assume the regular replication mode in this step.

During plasmid replication, pAMR may mutate into pAMRmut with probability $v$. For simplicity, we assume that at each cell division, at most one mutation occurs. The pAMRmut variant imposes a small fitness cost on the host cell, which increases linearly with the number of pAMRmut copies – up to a maximum factor $s$ (with $s$ being negative). Consequently, cells with $j$ pAMRmut copies have a reduced replication rate, scaled by the factor $1 + \frac{j}{n}s$.

Over time, the frequency of cells with pAMRmut copies will evolve towards mutation–selection balance, resulting in their low prevalence by the time pCRISPR is introduced in the subsequent step.

For a more detailed explanation and analysis of this step see Santer and Uecker (2020).

***Step 2: pCRISPR introduction.*** At time $t = 0$, pCRISPR is introduced to the resistant population in such a way that each cell receives exactly one pCRISPR copy. Once delivered, pCRISPR immediately exerts its interference activity. We distinguish two types of pCRISPR based on their mode of target interference:

**pCleaving:** In cells receiving pCleaving, all copies of pAMR are cleaved and lost before the next cell division. In contrast, pAMRmut copies, being protected by their mutation, remain intact and replicate according to their replication mechanism.

**pSilencing:** With pSilencing, the pAMR copies remain in the cell, but the expression of the AMR gene is silenced, so they no longer confer antibiotic resistance. Meanwhile, pAMRmut is not affected by this repression and continues to provide resistance. Importantly, the replication of both pAMR and pAMRmut proceeds normally under pSilencing. If, in later generations, all pSilencing copies are lost from a cell, the pAMR copies regain functionality and start providing resistance again.

If the introduction of pCRISPR causes the copy number $n$ to be exceeded (which can only occur in the case of incompatible pSilencing), this state persists until the next cell division, at which point only $n - 1$ plasmid replications occur.

***Step 3: Birth-death process after pCRISPR and antibiotic introduction.*** After antibiotics are added, we simulate the dynamics of the bacterial population using a birth-death process similar to the approach in Santer and Uecker (2020). Each cell divides and dies at rates that depend on the total number of pAMRmut and non-silenced pAMR copies within the cell, denoted as $k \in \{0, \dots, n\}$. This yields a total of $n+1$ cell types with different birth rates $\lambda_{\min}^{(n)} = \lambda_0^{(n)}, \dots, \lambda_k^{(n)}, \dots, \lambda_n^{(n)} = \lambda_{\max}^{(n)}$ and death rates $\mu_k^{(n)}$, which are defined analogously. During replication of pAMR, mutations from pAMR to pAMRmut may happen with probability $v$, where we make the simplifying assumption that cells that acquire the mutation *de novo*, carry it in only one copy before division.

Since the extinction probability depends solely on the ratio of birth to death rates, we compare results for different birth-death ratios $R_k^{(n)} := \lambda_k^{(n)}/\mu_k^{(n)}$, which we call the fitness of a cell. For cells with $k \in \{1,\ldots,n-1\}$, we assume that for the same copy number $n$ more functional AMR plasmids provide a higher or equal fitness, i.e. $R_k^{(n)} \le R_{k+1}^{(n)}$. Furthermore, when comparing different copy numbers, the maximum cell fitness is the same regardless of the specific value of $n$. In addition, we assume that pAMRmut confers the same level of resistance as non-silenced pAMR.

These considerations lead us to examine three fitness functions:

***Linear* fitness:** Fitness increases linearly with the number of functional AMR plasmids, i.e. $R_k^{(n)} = R_0^{(n)} + k/n * \left(R_n^{(n)} - R_0^{(n)}\right)$.

***Dominant* AMR plasmids:** A single functional AMR plasmid is sufficient for a cell to achieve maximum fitness, meaning that $R_k^{(n)} = R_n^{(n)}$ for $1 \le k \le n$.

***Recessive* AMR plasmids:** A cell requires all plasmid copies to be functional to benefit, so for $k < n$, it holds that $R_k^{(n)} = R_0^{(n)}$.

We do not factor in a cost of the total plasmid load, which may differ between cells if pAMR/pAMRmut and pCRISPR are compatible.

### Supplementary Note 2: Model evaluation and validation

This note provides a structured model evaluation and validation framework, loosely following the approach of Augusiak *et al.* (2014) and adapting it to our specific scenario.

**Parameter assumptions.** Our model includes four one-dimensional parameters, i.e. $N$, $n$, $s$, and $v$ and two $n+1$-dimensional parameters, i.e. the birth and death rates (see Table 1). For Figures 2, 3, and 5 exemplary parameters from a wide range have been chosen.

We use population sizes $N$ between $10^6$ and $10^8$, which is in line with the bacterial load of *E. coli* and *S. aureus* in the bloodstream of infected patients reported by Turkeltaub *et al.* (2024). For the plasmid copy number $n$, we consider values between 1 and 12, representing low to moderate copy number plasmids (Ramiro-Martínez *et al.* 2024), which are typically those encoding antibiotic resistance (Carattoli 2009). The mutation cost $s$ ranges from 0.05 to 0.8, where $s = 0$ corresponds to a cost-free mutation and $s = 1$ to lethal mutation, thus covering a broad spectrum of biologically relevant fitness costs. For the mutation rate $v$ we use a broad range of values between $10^{-7}$ and $10^{-9}$, as experimentally measured in a setting similar to ours by Li *et al.* (2023a), where multiple target loci and higher Cas expression were used to decrease the effective escape mutation rate. However, estimating mutation rates in a specific treatment setting is challenging, as this depends on numerous biological factors. Finally, birth and death rates in pathogenic populations vary greatly and can fluctuate during treatment (Grant *et al.* 2008; Kaiser *et al.* 2013). When combined, the birth and death rates used in our Figures result in a population doubling time of fully resistant cells and cells in the absence of treatment ranging from 2.4 to 15 generations, where one generation corresponds to the generations in the discrete process induced by our continuous time birth and death process. If one generation corresponds to a time span of the average time until a cell divides numbers are slighly lower.

While we numerically explored the effect of parameter variations (Fig. 5), we only considered a limited set of parameters. Therefore, we used analytical arguments to show that our key conclusion regarding the existence of a copy number threshold for the best pCRISPR variant remains valid for any parameter set. These arguments assume that the population size $N$ is sufficiently large and the mutation rate $v$ remains close enough to 0 (see Fig. 4 and Supplementary Note 3).

**Conceptual Model Evaluation.** Our model is based on a Moran process and a birth-death process, inheriting both the advantages and limitations of these approaches. For example, bacterial birth and death rates in natural populations are governed by much more complex regulatory mechanisms and environmental factors than those captured by a simple birth-death process.

Regarding the plasmid dynamics, we assume that plasmid replication happens immediately before cell division and that the plasmids are afterwards distributed equally to the two daughter cells.plasmid segregation SI While this is mathematically convenient and implements the fundamentally important stochasticity of plasmid inheritance, real plasmid inheritance mechanisms are less perfect. The number of plasmids given to each daughter cell is not necessarily equal, and plasmid copy numbers can vary between different cells (Shao *et al.* 2021). Moreover, plasmids can be replicated after (and not right before) they are distributed to the daughter cells Nordström (2006). However, for a similar model on the evolutionary dynamics of alleles on multicopy dynamics in a setting without CRISPR plasmids, theoretical predictions are in good agreement with experimental data (Garoña *et al.* 2023).

Furthermore, our model does not investigate pCRISPR delivery and delivery efficiency. However, in both, *in vitro* Hamilton *et al.* (2019) and an *in vivo* mouse models Neil *et al.* (2021), CRISPR-Cas tools could be delivered to 100% of the target population. This, together with the fact that the process of conjugation results in exactly one plasmid copy being delivered to a recipient cell, makes the model assumption of each cell receiving one plasmid at the beginning of the treatment reasonable.

Next, we assume that CRISPR-Cas interference is immediate and cleaves all target plasmids present in the cell simultaneously. Experiments for copy numbers up to 70 (Citorik *et al.* 2014) and 100–300 (Tagliaferri *et al.* 2020) have shown that all plasmid copies can be cut. An interesting option here is that CRISPR-Cas proteins fused to a relaxase could be transferred, which entirely eliminates expression time (Guzmán-Herrador *et al.* 2024).

Lastly, our model does not account for potentially important ecological factors, such as horizontal plasmid transfer, spatial structure, resource competition, pharmacokinetically driven fluctuations in antibiotic concentrations during treatment, and the delivery efficiency of pCRISPR. We acknowledge that these factors could influence dynamics of plasmid and bacterial populations and consider this in the discussion. However, it is important to mention here that our model is not intended to directly inform patient treatment. It could also be applied to controlled environments, such as *ex vivo* settings, where bacterial populations are treated outside the host.

**Model output verification.** The key result of this work is the differential effect of pCleaving and pSilencing on sensitizing bacteria, with respect to the plasmid copy number. The two strategies are qualitatively different because for almost all considered parameters there is a threshold copy number above which pSilencing is superior (shown in Supplementary Note 3).

While most research has focused on cleaving CRISPR-Cas, recent studies like Yao *et al.* (2022) and Mayorga-Ramos *et al.*

(2023) have begun to explore the effectiveness of silencing mechanisms as well. However, currently there is to our knowledge no experimental data available to directly compare the effectiveness of cleaving and silencing systems.

By introducing incompatible and compatible cleaving and silencing CRISPR-Cas systems against low- and high-copy AMR plasmids and comparing the efficacy against each other, future studies could empirically validate our findings, especially as the steps of delivery and CRISPR-Cas activity can be entirely separated in *in vitro* experiments.

### Supplementary Note 3: Existence of threshold copy number

The goal of this note is to determine whether a critical copy number exists at which incompatible pCleaving is surpassed by compatible pSilencing as the optimal method. Note that we argue mainly heuristically in this note and it should thus not be considered to be a rigorous proof.

As discussed in the Results section, incompatible pCleaving consistently outperforms compatible pSilencing for copy numbers $n = 1$ and $n = 2$, but compatible pSilencing may or may not become more effective for sufficiently high copy numbers.

A higher copy number has two effects, it changes the standing genetic variation at the start of the treatment and increases the number of plasmids that can acquire *de novo* mutations.

**Copy number affects the SGV at the start of the treatment.** We again consider $f^{(\mathrm{SGV})}$, the equilibrium distribution of pAMR and pAMRmut copies, prior to pCRISPR introduction and antibiotic treatment. When the copy number $n$ increases there is a linear increase in the emergence of novel mutations within wild-type cells, but it becomes increasingly difficult for cells to reach the state in which all pAMR copies have been replaced by pAMRmut copies. To make this more quantitative, let us consider a cell that has just acquired a pAMRmut through mutation. In a population of constant size, the cell lineage would on average remain at one cell if the process were neutral. In such a neutral scenario, the mutation would have a probability of $1/n$ of fixing in the cell lineage. Since pAMRmut carries a cost, the probability that a mutant homozygous cell appears is less than $1/n$. Thus, despite the mutational input that linearly increases with $n$, the frequency of homozygous mutant cells decreases as $n$ increases, as illustrated in Fig. S5. For large copy numbers, the population thus consists predominantly of heterozygous cells with a small fraction of pAMRmut copies. Immediately after the introduction of pCRISPR into the population, incompatible pCleaving then results in a substantial number of cells with a high fraction of (replicated) pAMRmut copies. Conversely, compatible pSilencing does not alter the cell types from the SGV, as illustrated in Fig. 2, meaning most cells still contain a very low amount of pAMRmut copies.

Let us now consider how this changes the extinction probabilities.

**Linear fitness function.** If the cell fitness increases linearly with the number of pAMRmut copies during antibiotic treatment, cells with a higher number of pAMRmut copies have a substantially lower extinction probability than those with few copies. During antibiotic treatment, the establishment probabilities of a cell with a given number of functional pAMRmut copies is the same in both CRISPR-Cas treatment scenarios: in one case, pAMRmut competes with pCRISPR, in the other with silenced pAMRmut.

As $n$ approaches infinity, the fraction of WT cells in the SGV declines to 0. Consequently, after pCRISPR introduction, the population consists of (in comparison) many cells of high fitness for incompatible pCleaving and a population with few cells of high fitness for compatible pSilencing. Since the impact of *de novo* mutations on the extinction probability of the population becomes negligible compared to the number of pAMRmut copies reproduced after pCRISPR introduction, for sufficiently high copy numbers $n$, compatible pSilencing is more likely to drive the population to extinction than incompatible pCleaving. (This heuristic argument assumes that the maximal fitness benefit is constant and does not grow with the copy number, which we assume throughout the manuscript.)

For *dominant* and *recessive* fitness models, the argument above cannot be used, as, after pCRISPR introduction, the fitness of cells with few and many pAMRmut is the same. Therefore, a different approach is needed.

**Recessive fitness function.** Let $f^{(\mathrm{pCleaving})}$ and $f^{(\mathrm{pSilencing})}$ be the frequencies of the different cell types one generation after pCRISPR has been introduced into the cells and took effect. For the recessive fitness function the only chance for a cell lineage to escape extinction is to evolve into a homozygous pAMRmut cell. The probability of doing so increases the more pAMRmut copies the initial cell has. As the frequency of homozygous mutant cells in the SGV decreases, as $n$ increases, but, unlike pSilencing, pCleaving increases the number of pAMRmut copies after its introduction, this leads for *recessive* fitness to

$$\frac{P_{\mathrm{ext}}^{(\mathrm{SGV\,pCleaving})}}{P_{\mathrm{ext}}^{(\mathrm{SGV\,pSilencing})}} := \frac{\prod_{j=1}^{n} p_{\mathrm{ext}}(0, j, n-j)^{f^{(\mathrm{pCleaving})}(j)}}{\prod_{j=1}^{n} p_{\mathrm{ext}}(n-j, j, 1)^{f^{(\mathrm{pSilencing})}(j)}} \overset{n\to\infty}{\longrightarrow} 0\,, \tag{5}$$

using the same notation as in the main text. However, this analysis does not account for *de novo* mutations, which could considerably lower the effectiveness of compatible pSilencing. To assess the probability of evolutionary rescue due to these *de novo* mutations in the case of a *recessive* fitness function, we introduce the random variable $X$, which represents the number of times cell lineages originating from a single mother cell that just acquired a *de novo* mutation evolved into a homozygous mutant cell lineage. For a recessive fitness function only those cell lineages have the chance to survive. For a *recessive* fitness function, the expectancy of $X$ must be lower than $1/n$, as only heterozygous cells are considered and their expected 'number of offspring' is lower than 1: If the number of offspring were exactly one, the fixation probability of the pAMRmut copy would be exactly

$1/n$ (as for any plasmid), but since a heterozygous cell under treatment has on average fewer than one daughter cell until it dies, the probability of the one pAMRmut copy establishing itself in a cell lineage is less than $1/n$. Jensen's inequality theorem then leads to

$$p_{\text{ext}}(n-1,1,1) = E\left(p_{\text{ext}}(0,n,1)^X\right) = E\left(R_{\text{max}}^{-X}\right) \geq R_{\text{max}}^{-E(X)} \geq R_{\text{max}}^{-\frac{1}{n}}. \tag{6}$$

since $h(x) := a^x$ is a convex function for $a > 0$. Following Eq. (4) of the main part, we now get

$$P_{\text{ext}}^{(de\ novo)} \approx e^{-v\,n\,N\,\frac{R_{\text{min}}}{1-R_{\text{min}}}(1-p_{\text{ext}}(n-1,1,1))} \geq\ p_{\text{ext}}(n-1,1,1)^{v\,n\,N\,\frac{R_{\text{min}}}{1-R_{\text{min}}}} \geq R_{\text{max}}^{-v\,N\,\frac{R_{\text{min}}}{1-R_{\text{min}}}}\,, \tag{7}$$

as $e^{-(1-x)} \geq x$ for $x \in [0,1]$. This means that $P_{\text{ext}}^{(de\ novo)}$ does not approach zero as $n$ increases, as the right-hand side of the equation is not dependent on $n$. For the total extinction probabilities $P_{\text{ext}}^{(\text{total pSilencing})} := P_{\text{ext}}^{(de\ novo)}\, P_{\text{ext}}^{(\text{SGV pSilencing})}$ and $P_{\text{ext}}^{(\text{total pCleaving})} := P_{\text{ext}}^{(\text{SGV pCleaving})}$ as in Eq. (1) of the main part, combining Eq. (5) and Eq. (7) then leads to

$$\frac{P_{\text{ext}}^{(\text{total pCleaving})}}{P_{\text{ext}}^{(\text{total pSilencing})}} := \frac{1}{P_{\text{ext}}^{(de\ novo)}}\,\frac{P_{\text{ext}}^{(\text{SGV pCleaving})}}{P_{\text{ext}}^{(\text{SGV pSilencing})}} \overset{n\,\to\,\infty}{\longrightarrow} 0\,. \tag{8}$$

Therefore, for *recessive* fitness, there must always exist a threshold copy number so that compatible pSilencing is superior above that threshold.

This leaves the *dominant* fitness function as the last variant to consider.

**Dominant fitness function.** Here, the threshold copy number does not always exist, and incompatible pCleaving can outperform compatible pSilencing for arbitrarily large copy numbers, for example when $R_{\text{max}}$ is relatively high. For instance, a threshold does not exist for parameters $R_{\text{max}} = 2.3$, $R_{\text{min}} = 0.95$, $v = 10^{-8}$, $s = -0.1$, $N = 10^7$.

### Supplementary Note 4: Figures for *regular* replication

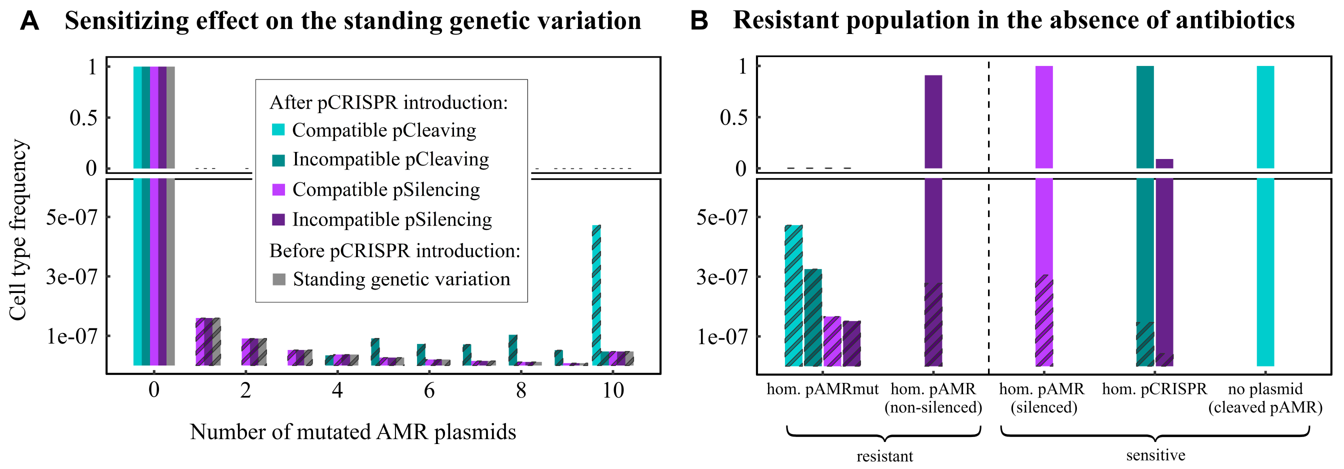


**Supplementary Fig. S1. The standing genetic variation (SGV) before and after the introduction of CRISPR-Cas to the population for copy number $n = 10$ (A) and the long-term evolution of the cells carrying at least one mutation without considering *de novo* mutations and in the absence of antibiotics (B) for *regular* replication.** The SGV before the introduction of pCRISPR to the population for copy number $n = 10$ is depicted in grey in A. It illustrates the frequency of cells carrying 0 to 10 pAMRmut copies, while the other plasmids inside the cell are WT pAMR copies. The other bars show the frequency of all cell types after the different pCRISPR copies are introduced. Parameters: $v = 10^{-8}$, $n = 10$, $s = -0.1$.

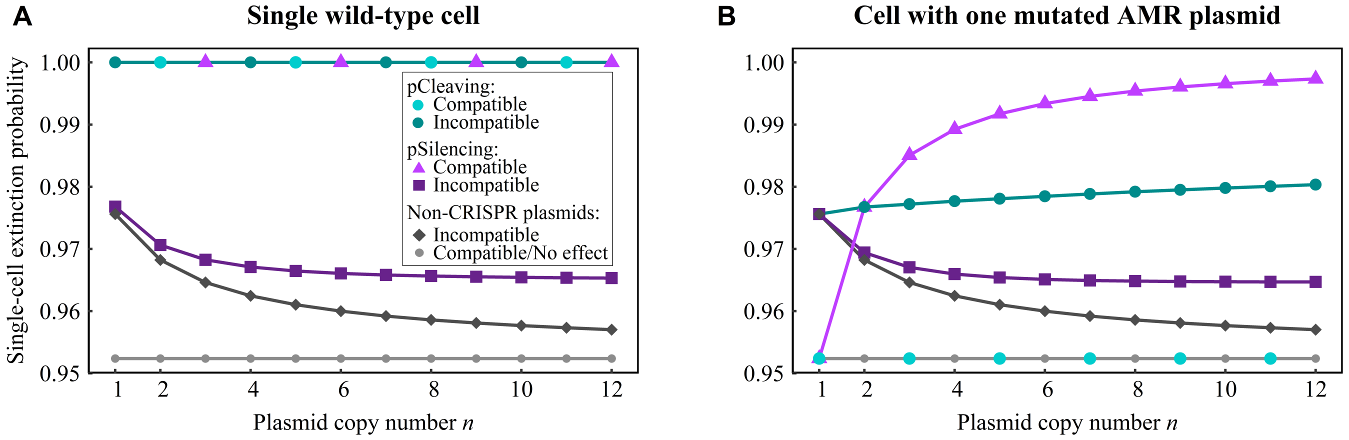


**Supplementary Fig. S2. Extinction probabilities for a single cell lineage due to antibiotic treatment after pCRISPR introduction for *regular* replication.** The long-term extinction probability for offspring derived from single wild-type cells (A) and cells carrying exactly one pAMRmut and $n-1$ WT pAMR copies (B) prior to the CRISPR-mediated cleaving or silencing. For comparison, plasmids without a CRISPR-Cas system are also shown. Since a compatible non-CRISPR plasmid has no effect, this scenario is equivalent to the antibiotic treatment of a resistant population. The analysis uses a linear fitness function with parameters: $R_{\max} = 1.05$, $R_{\min} = 0.95$.

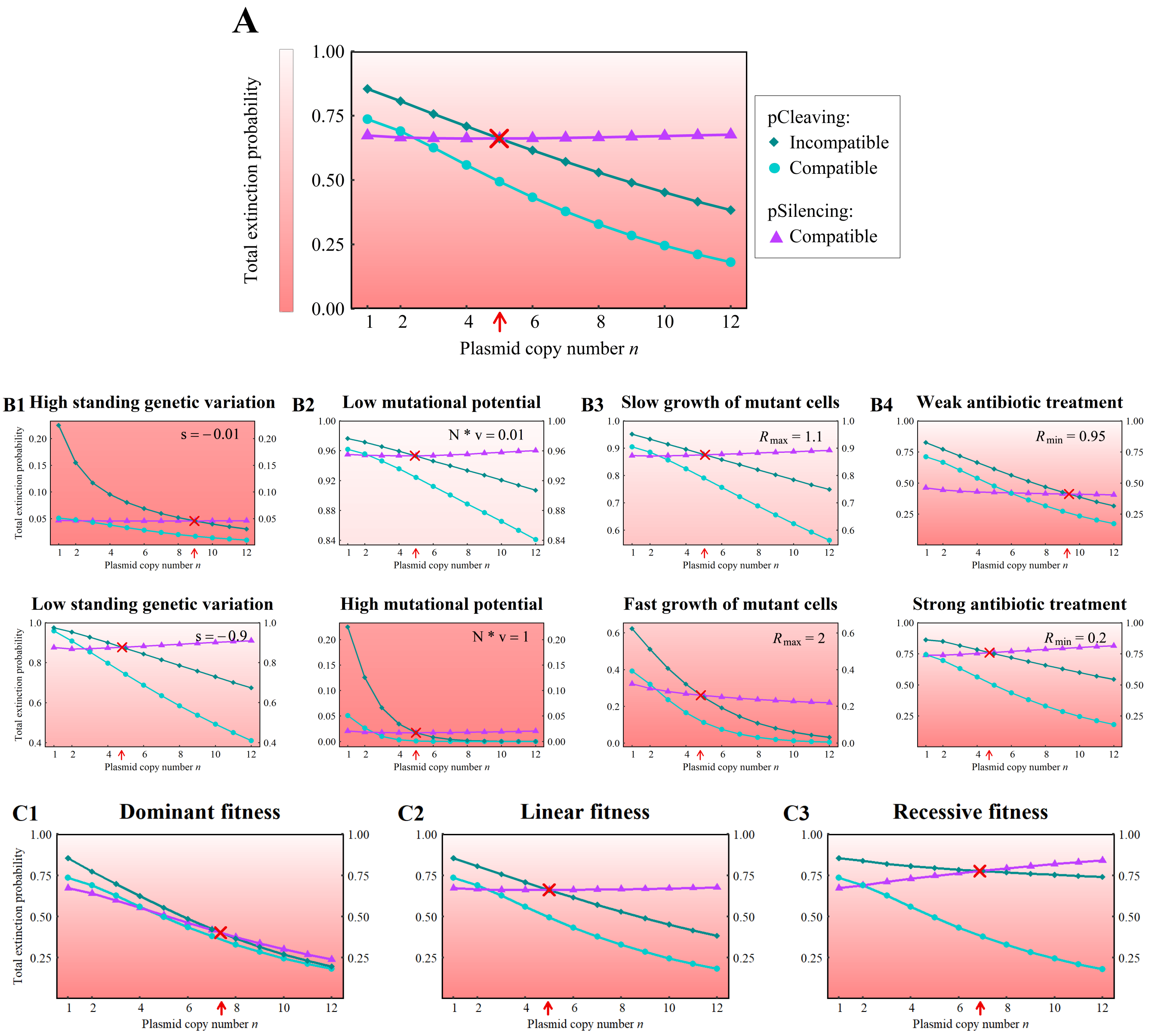


**Supplementary Fig. S3. Total extinction probability in a pathogenic population for *regular* replication.** The focus is only on the following three primary cases: compatible and incompatible pCleaving, and compatible pSilencing. (A) shows a typical behavior of the three curves. The chosen parameters are $R_{\max} = 1.3$, $R_{\min} = 0.8$, $v = 10^{-8}$, $s = -0.1$, $N = 10^7$. The fitness function used is linear. (B) illustrates the effect on (A) when a specific parameter is adjusted to a higher or lower value as indicated in the top right-hand corner of each plot. All other parameters remain constant as in (A). In (C), the fitness function is varied while the other parameters are the same as in (A). The red arrow and cross indicate the point of intersection of the two curves, representing compatible pSilencing and incompatible pCleaving. Below this intersection point, incompatible pCleaving is consistently superior, while above it, compatible pSilencing emerges as the more effective strategy. Note the different ranges on the y-axis across panels. The gradient of the background color from white to light red visually represents the success of the treatment.

### Supplementary Note 5: Additional Figures

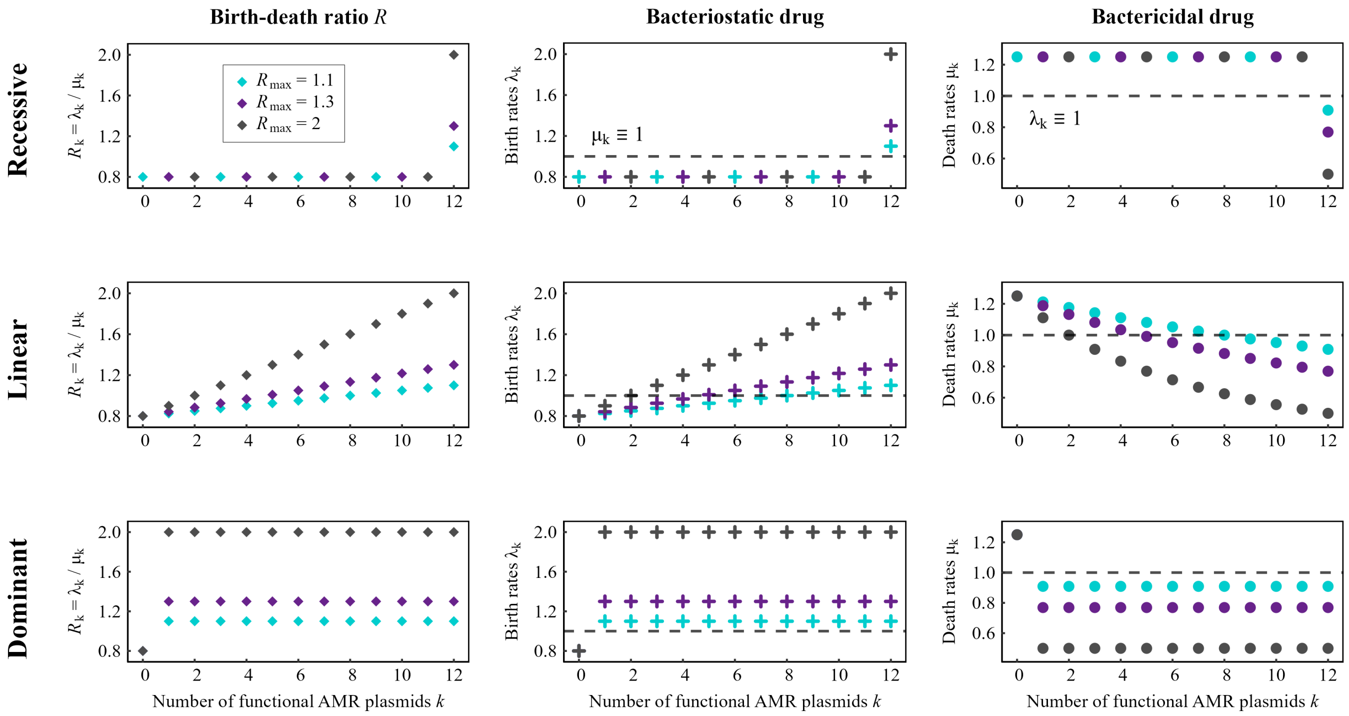


**Supplementary Fig. S4. Birth and death rates corresponding to the fitness/birth-death ratios used in Fig. 5 for *dominant*, *linear*, and *recessive* fitness functions under bacteriostatic and bactericidal treatments.** In the bacteriostatic scenario, antibiotic action reduces the birth rate $\lambda_k$ of non-homozygote mutated cells (i.e., not fully resistant, $k < n$), while the death rate $\mu_k$ remains constant regardless of the level of resistance. Conversely, under bactericidal treatment, the antibiotic increases the death rate $\mu_k$, while the birth rate $\lambda_k$ remains constant across all cell types (i.e. varying numbers of functional AMR plasmids). Parameters: $n = 12$, $R_{\min} = 0.8$.

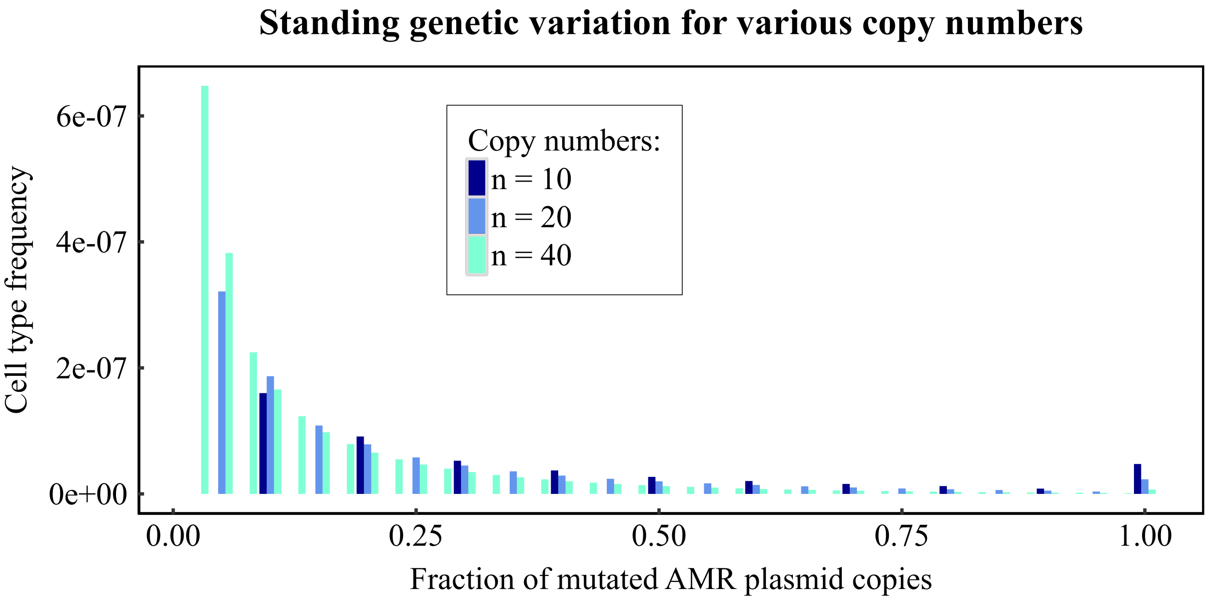


**Supplementary Fig. S5. The standing genetic variation (SGV) prior to pCRISPR introduction for different copy numbers:** With increasing copy number, the frequency of cells that carry at least one pAMRmut copy increases as more WT pAMR copies can mutate. In contrast, it becomes increasingly difficult for cells to arise that are homozygous for pAMRmut. In the theoretical scenario where $n \to \infty$, the probability of a cell lineage that originates from a wild-type cell to evolve into a homozygous mutant cell converges to 0. This can be easily seen considering a cell lineage that originates from a cell with a neutral mutation on a single plasmid. The cell lineage will either evolve into a homozygous mutant cell with probability $1/n$ or lose the mutated plasmid. In the case of pAMRmut with a small fitness cost the probability is smaller than $1/n$. Consequently, as the copy number increases, the frequency of homozygous mutant cells in the SGV decreases. Parameters: $v = 10^{-8}$, $s = -0.1$.

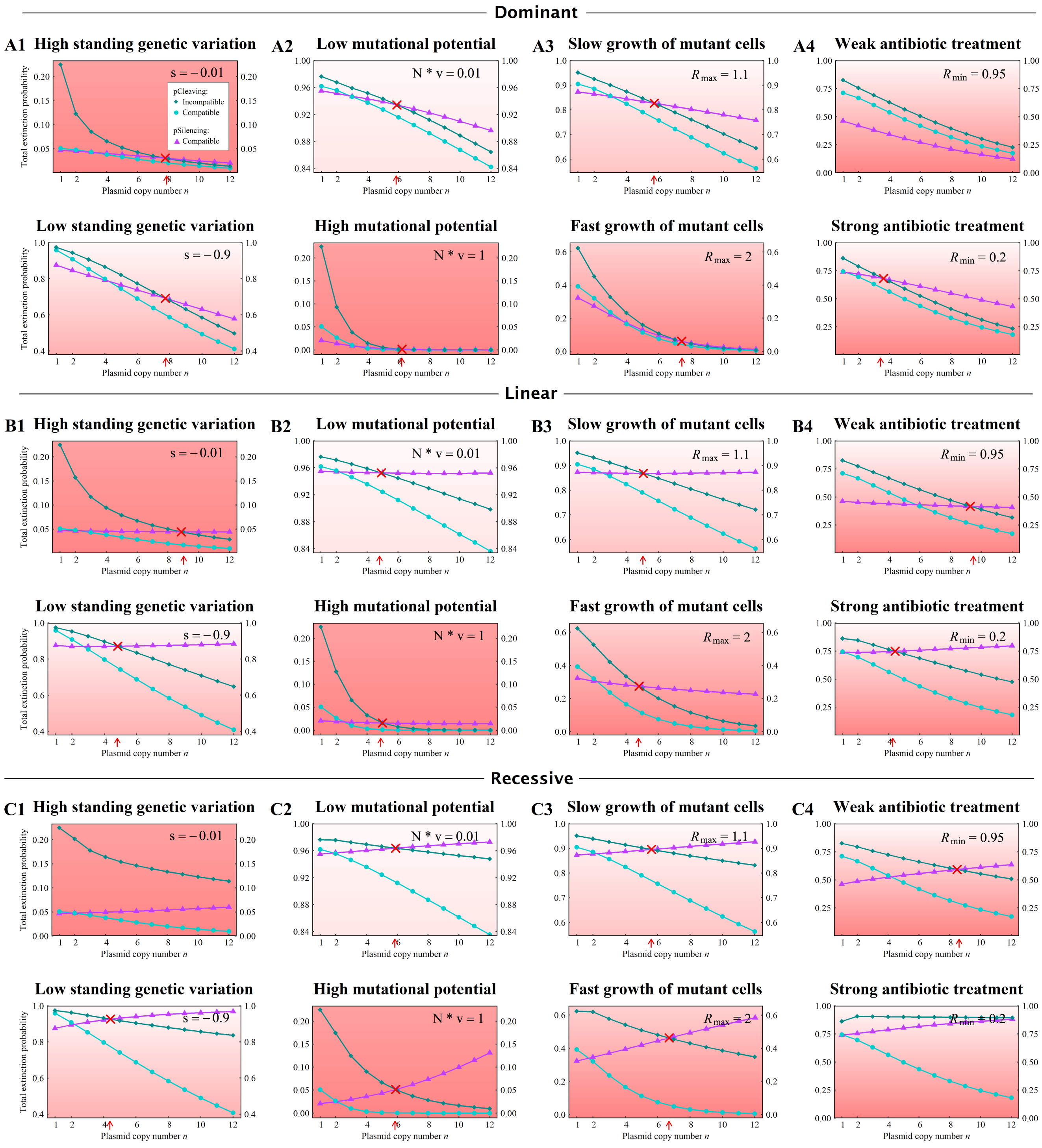


**Supplementary Fig. S6. Population extinction for *dominant* (A1 - A4), *linear* (B1 - B4) and *recessive* (C1 - C4) fitness functions.** As in Fig. 5 B, the default values are $R_{\max} = 1.3$, $R_{\min} = 0.8$, $v = 10^{-8}$, $s = -0.1$, $N = 10^7$, $u = 0$, while one specific parameter is adjusted to a higher or lower value as indicated in the top right-hand corner. The red arrow and cross indicate the point of intersection of compatible pSilencing and incompatible pCleaving. Note the different ranges on the y-axis across panels. As can be seen, the fitness function can alter the monotonic trend of compatible pSilencing as well as the threshold copy number. If the fitness function is dominant, the threshold does not always exist.

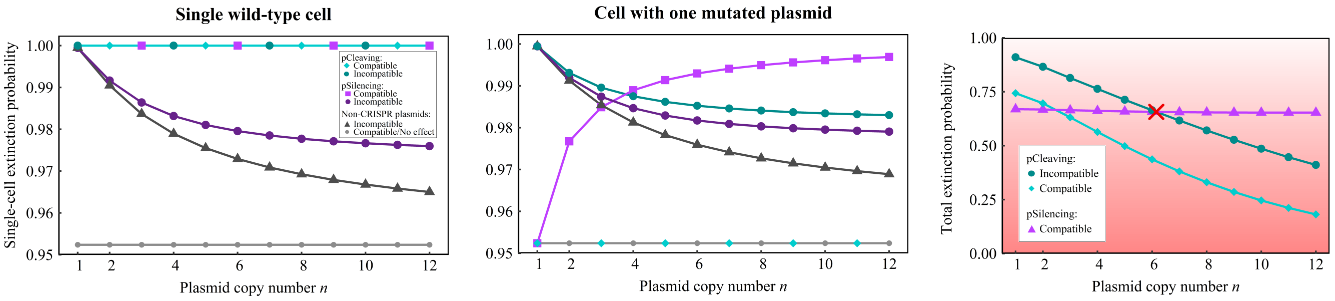


**Supplementary Fig. S7. Constant influx of CRISPR-Cas plasmids.** Results for a modified version of our model where, after acquiring a pCRISPR at $t = 0$, each cell in the population has the chance to receive additional pCRISPR copies afterwards (i.e. for $t > 0$) at rate $u$. Compared to receiving only one pCRISPR per cell, cells carrying mutations at $t = 0$ have a higher chance of going extinct if pCRISPR is incompatible. However, due to the same performance on WT cells, the extinction population of the whole population remains similar. Incompatible pSilencing is still not an effective strategy as plasmid losses due to segregational drift outweigh the influx of pCRISPR. Parameters: $u = 0.1$, $R_{\max} = 1.3$, $R_{\min} = 0.8$, $v = 10^{-8}$, $s = -0.1$, $N = 10^7$.

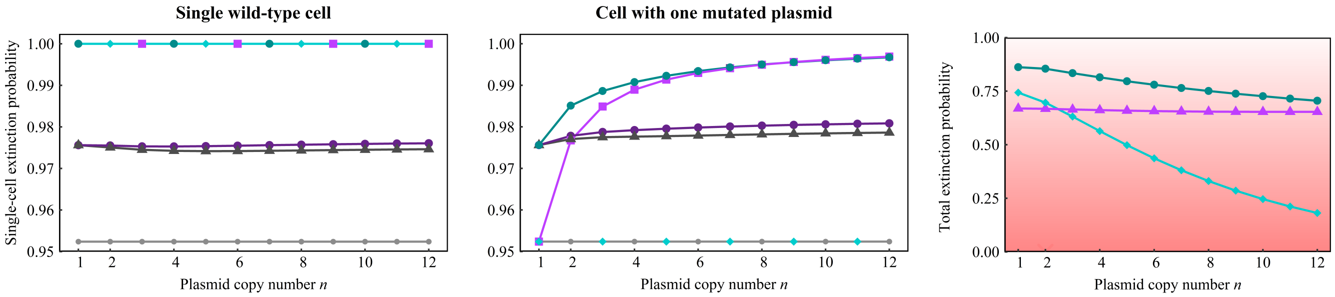


**Supplementary Fig. S8. Replication advantage of the CRISPR-Cas plasmid (twice as fast replication).** Results for a modified version of our model, where the replication rates of pCRISPR and pAMRmut differ. This figure shows the outcome where each of the pCRISPR copies is twice as likely to replicate as the pAMR or pAMRmut copies. Naturally, this scenario is only considered for the *random* replication model. While this change has no impact on compatible pSilencing and pCleaving in comparison to equal replication speed, it enhances the efficacy of incompatible pCleaving. Parameters: $R_{\max} = 1.3$, $R_{\min} = 0.8$, $v = 10^{-8}$, $s = -0.1$, $N = 10^7$.
